# Phenology and flower species availability define wild bee communities on river embankments

**DOI:** 10.1101/2025.05.16.653962

**Authors:** Constant Swinkels, Linde Slikboer, Ivo Raemakers, Niels Godijn, Eelke Jongejans, Hans de Kroon

## Abstract

River embankments cover extensive areas in estuaries worldwide, yet their potential for pollinator conservation remains largely unexplored. Even more so than road verges, these embankments may offer widespread habitat for pollinators due to their size and sun-facing warm slopes. Here we present a comprehensive analysis of bee communities from 157 dike grasslands along the lower branches of the Meuse and Rhine rivers in the Netherlands in relation to vegetation composition and landscape characteristics. We encountered nearly half (156) of all Dutch bee species. Almost 10% of all individuals belonged to red-listed species, especially in early spring when we observed rich ground nesting bee communities with their associated kleptoparasites. Wild bee abundance and diversity increased with more flowering plant species present, but saturated at low flower species richness. This early saturation suggests that even modest conservation efforts could effectively alleviate floral resource limitation for pollinators. In contrast to the overall pattern, red-listed species occurrence increased with flower diversity without saturating effects and correlated stronger with landscape level factors. Together, these results highlight the unique conservational value of river embankments for wild bees and illustrate the unused potential at many dike grasslands worldwide.

## 1. Introduction

Evidence of flower visiting insect decline in North-Western Europe is mounting [1–5]. These pollinators are crucial for ecosystem functioning through the pollination of wild flower species [6,7], and some species provide additional ecosystem services such as crop pollination [8,9]. Wild bees are amongst the most important pollinators, both for their efficiency [10] and ubiquity [11]. Suffering from a range of anthropogenic influences [12,13], decrease in habitat availability and quality with subsequent loss of floral resources is generally seen as the major contributor to this decline [1,5]. Consequently, restoration practices aim to restore high-quality habitats, primarily through the promotion of local floral resource abundance and diversity.

Although nature reserves provide suitable habitat, in Western Europe they are often small and isolated. Hence, it is essential to create habitat at the landscape level to promote wild bees [14]. In this context, linear landscape elements (LLE), such as road verges, railway lines, power lines, ditches and embankments, can contribute to bee conservation [15]. LLEs form an extensive network of semi-natural grassland across countries, interweaving natural, rural and urban areas [16]. Studies show that well-managed LLEs can promote both flowers and bees (e.g. [17–19]), including red listed and endangered species of specific conservation interest [18,20].

However, although the potential of LLEs for bee conservation is evident [15,21], we lack a broad overview of the communities they harbour and the conditions associated with community richness. This is particularly true for dike grasslands. In lowland regions, dikes can cover a considerable area, signified by the more than 17,000 kilometres of dikes found in the Netherlands [22]. Dikes are notably different from other LLEs for a number of reasons. By definition these embankments along water bodies are raised above the surrounding flat landscape, with warm, sun facing slopes as potential hotspots for bee nesting [23]. In addition, they are often larger in surface area and relatively old compared to many other linear landscape elements. Both grassland age and size have been related to bee community richness [24]. Finally, vegetation on dikes contributes to their flood protection services, with studies suggesting that forb-rich grasslands could benefit dike stability due to increased rooting densities [25]. With increasing efforts to extend the area of forb-rich dike communities for flood protection, dikes have a high potential to effectively contribute to bee conservation as additional benefit.

Here we present a comprehensive analysis of dike grassland bee communities along the lower branches of the Meuse and Rhine rivers in the Netherlands. We collected and analysed multi-year data on bee community and vegetation composition from 157 locations on dike grasslands. The aims of this study were (1) to characterise the current dike grassland wild bee community composition in relation to the wild bees of the Netherlands, (2) to analyse the relationship between wild bee species richness and abundance with local flower species richness in space and time, and (3) to examine how the presence of specific bee species relates to their food sources. Herewith, we reveal the conservation potential of river embankments for wild bees in a West-European lower estuary, and present relevant relationships between bee and wild plant communities informing wild bee conservation efforts.

## 2. Materials and Methods

### 2.1 Location

Samples were collected at 157 locations from dikes in the southern half of the Netherlands, essentially covering the entire Dutch lowland river and Delta area (figure 1). Data were collected from a range of different dike grasslands, including monotone grasslands devoid of flowers and grasslands with high flower abundance and more structural diversity including shrubs and small trees. The study locations include primary dikes, currently functioning as flood protection barrier, and secondary dikes, no longer in use as flood protection barrier. Primary dikes are nowadays constructed from heavy clay, often with a top soil layer of more sandy substrate, and are mown twice a year (June and September) when managed with the intent to facilitate biodiversity [26]. Many of these primary dikes were ‘renewed’ about 30 years ago after the highwaters of 1993 and 1995, meaning vegetation has had time to develop since. Secondary dike grasslands are not renewed (sometimes 100+ years old grasslands), with diverse soil types and subjected to a variety of management practices (e.g. infrequent mowing or grazing by cattle and sheep).

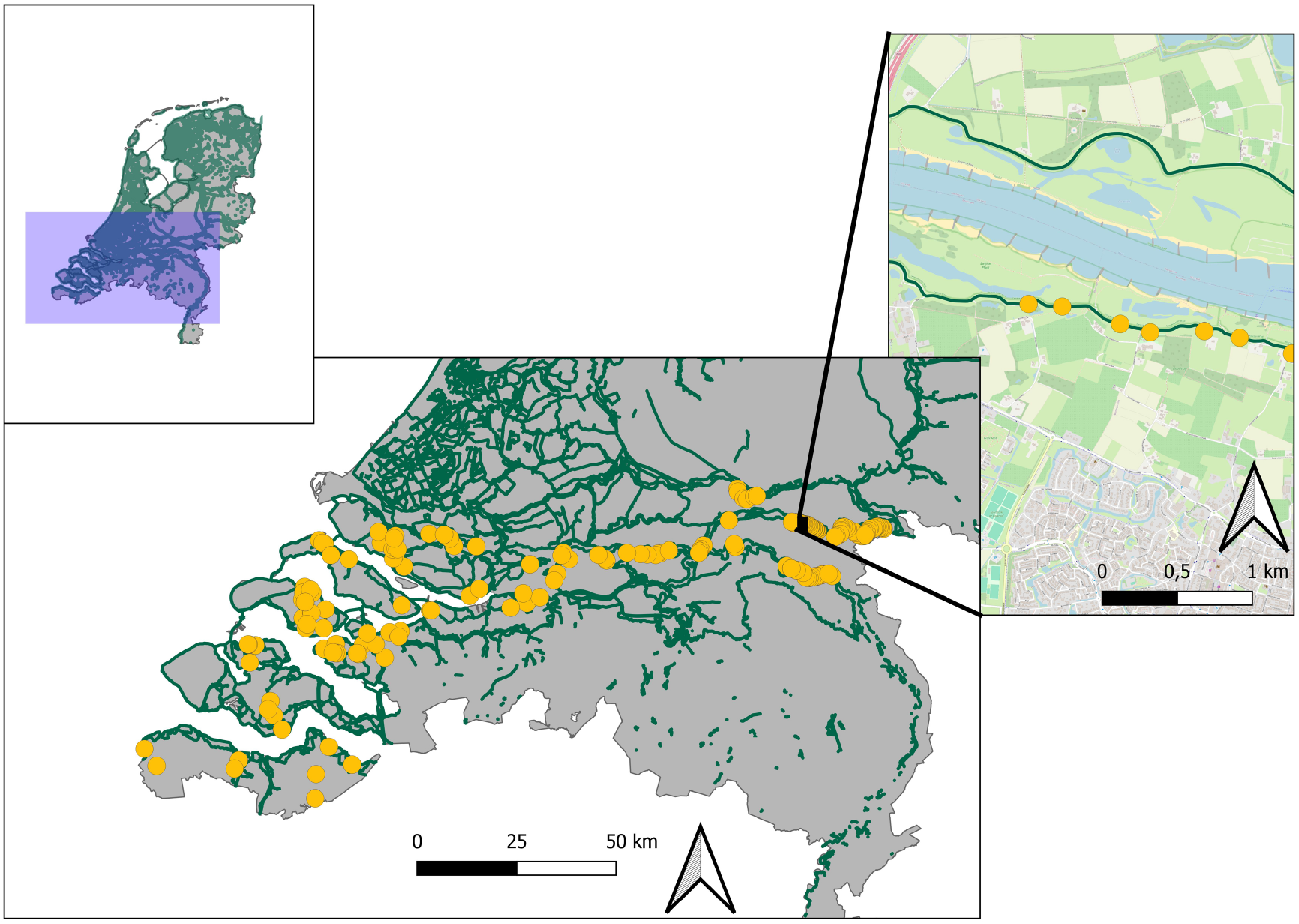

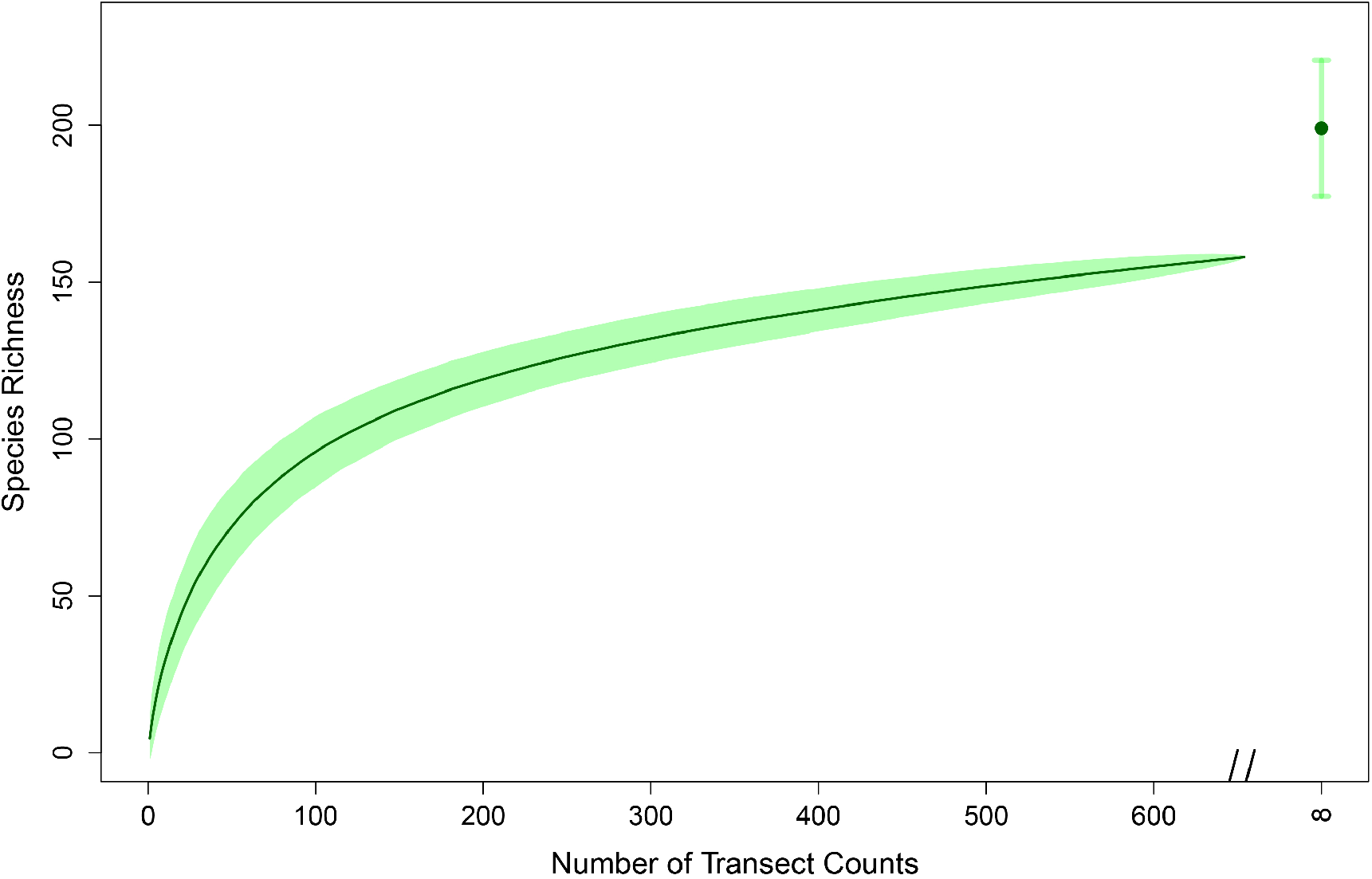

### 2.2 Sampling strategy

Samples were collected by experienced observers; Constant Swinkels (CS) in the east of the Netherlands, Ivo Raemakers (IR) in central and east NL, and Linde Slikboer and Niels Godijn (LS) in the South-Western delta. All observers visited locations multiple times per year to account for phenology of bees and plants. Data were collected from 2018 up until 2022 and in total 654 transects surveys were collected from the 157 locations, with a total sampling time of 181.5 hours.

#### 2.2.1 Bee Sampling

Bees were sampled along 150m long transects, running through representative vegetation for that dike grassland. Transects were sampled for 15 minutes of active searching time (time was paused while tracking, handling and identifying individuals) and all bees found within 1.5m on either side of the transect were recorded. Species that could not be identified in the field were collected and transferred to the lab for identification. The *Bombus terrestris* species complex was treated as one species due to the difficulties involved in correct identification of these species [27]. Honeybees were counted, but not included in further analysis.

For 81 transects, sampling time was longer than 15 minutes. For these, we corrected for sampling effort by randomly removing a percentage of individuals equal to the percentual time increase, assuming that bee individuals were evenly distributed over the transect. Individuals were removed stepwise, with chances of removal being equal for each individual. Species were eliminated from the sample if their abundance reached zero.

After identification, we recorded information on the status of species from the Dutch red list of bees [28]. In addition, we recorded species pollen preference from the German bee atlas [29] supplemented with information from the Dutch field guide on bees [30].

#### 2.2.2 Vegetation sampling

For each transect count local flower species richness and abundance was recorded, with all open flowers identified to the species level. Grass inflorescences were not included as these are not used by bees in our study area. The exact survey method differed for observers. CS counted all flowering plants per species within a 5×5m quadrat, IR counted floral units per species along the 150m netting transect (see bee sampling) and LS scored flower abundance on the Tansley scale. As such, the methods give comparable flower species richness (which is known to be a good predictor for bee diversity [31]), but flower abundance could not be compared between datasets (we present analysis along a flower abundance gradient per observer in appendix A.1 – A.3, finding similar patterns as our main analysis). For 33 transects flower data were missing, resulting in their exclusion from model analysis.

#### 2.2.3 Environmental factors

We collected information on landscape elements in a 250m and 1000m radius surrounding each location from the open access Dutch national Basic Registration Topography database. Land use types were divided in broad categories; agricultural area, semi-natural grassland, forest, fruit crops and a rest group (water, roads etc). Specific land use types within each category (e.g. crop type) are expected to influence bee communities in largely similar ways. Next to this we calculated the area of ‘NatuurNetwerk Nederland’ (NNN), designated natural area, in a 250m and 1000m radius and the distance to the closest NNN area. We collected weather data from all 50 Dutch weather stations from the KNMI database and interpolated measurements to our locations, where measurement weight was scaled inversely with distance. We collected information on daily average temperature, daily average windspeed, total daily rain and the percentage of sunshine during daylight hours.

### 2.3 Analysis

All data were analysed in R version 4.2.2 [32]. For analysis and graphics, packages lme4 [33] and ggplot2 [34] were used. For each section we describe what data we used, a flowchart can be found in Appendix B.1.

#### 2.3.1 Community composition

Using all available data, we calculated median number of individuals encountered per transect, that we deemed more informative than the average since abundance data was negative binomially distributed (Appendix B.2). As species abundances were skewed with few abundant species and many species occurring in low numbers (Appendix C.1), a common occurrence in wild bee communities [35], we identified a ‘core community’ consisting of species detected on at least 15 out of 157 locations (∼10% of all locations). A species accumulation curve was constructed for visual analysis by taking the total number of unique species encountered with increasing number of monitoring surveys. We randomly reordered surveys 1000 times, after which the mean curve and 95% confidence interval was plotted. We extrapolated species richness to estimate total species richness using the Chao estimator [36,37], as it has been found to perform best among competing estimators [38].

#### 2.3.2 Activity patterns

To study when different groups of bees most depended on dikes, we visually inspected activity patterns for the most abundant genera (*Andrena, Nomada, Bombus*, and a combination of the closely related *Lasioglossum* and *Halictus*, both Halictidae) and a group including all other genera (which separately had too few observations to produce meaningful patterns). The number of individuals observed within each genus for each transect count, including zeros, were plotted on a log scale. For the analysis of red listed species activity, Halictidae were added to the ‘other’ genera as they did not include many red listed individuals. We fitted a loess curve smoother to aid visual interpretation of the plots.

#### 2.3.3 Bee abundance and diversity

For all abundance and species richness analyses, four different (sub)sets of the data were used. Firstly, we included all available data. In addition, we split our dataset between bumblebees and all other wild bees. Bumblebees are eusocial and live in colonies, have large foraging ranges and are highly generalist. Their responses to local vegetation are well studied, with articles showing strong relationships between increases in flower availability and bumblebee abundance (e.g. Carvell et al., 2007; Pywell et al., 2006). Other wild bee responses, however, are relatively less studied, while they make up for most bee diversity and often have specific resource and nesting requirements. Finally, we selected only red-listed species to study what factors steered the occurrence of vulnerable species in our dataset.

We analysed bee abundance (negative binomial distribution, AppendixB.2) and species richness (Poisson distributed) using Generalised Linear Mixed Effect Models (GLMMs). Explanatory variables were selected based on ecological relevance. We include local flower species richness as this was the primary interest of our study. We studied phenological patterns by including the Julian date. We accounted for weather effects by including the percentage hours of sunshine during the day (calculated from the maximum possible hours of sunshine given the date) as most other weather variables are correlated with Julian date. As we repeated surveys at locations, sampling site was included as random factor. Finally, we included one of seven different landscape level factors; area covered by agriculture in a 250m and 1km radius, grassland in 250m and 1km radius, area of NatuurNetwerk Nederland (NNN) in a 250m and 1km radius and distance to the closest NNN area. NNN is a network of areas designated for nature protection in the Netherland. In addition, we ran a model without any landscape factors added. All explanatory variables were standardised (scaled) to allow comparison of model parameter estimates.

Initial models did not converge when including observer/region effect. To check whether including this effect could be expected to change model outcome, we compared model residuals for observers using a Kruskal-Wallis test. Only for models on bumblebee abundance and species richness we observed a significant relationship between the model residuals and observer/region (appendices D.1 and D.2), indicating dropping observer/region in these models was not justified. To study whether accounting for observer/region would change model outcomes, we constructed an alternative GLM including observers as fixed factor and dropping the location random factor, that explained limited variance (<0.1) compared to other fixed factors (appendices D.3-D.8), so that models converged.

For each model, we analysed different polynomials for flower species richness to determine what curve best described bee abundance and diversity along a flower-richness gradient. As overall third polynomial models were most parsimonious (appendices D.9 and D.10) we report significance values and model predictions from third polynomials in the results. As models described non-linear relationships, we performed additional breakpoint analysis on the simple linear relationship between bee abundance/richness and flower richness to find support for breakpoint values from visual inspection of the models.

Red-listed species abundance data were zero-inflated with few high abundance values (when colonies were present) and red-listed species richness was low. As such, we present presence/absence data (binomial) for red-listed individuals, constructing a GLM further using the same approach as described above. For all models we report marginal pseudo-R^2^ following [41].

#### 2.3.4 Predicting species occurrence based on resource preference

To analyse whether resource availability (i.e. presence of suitable flower host species as pollen source) could predict the occurrence of specific species, species specific resource preferences (see under 2.2.1) were matched with available flower species at each location to indicate the potential number of resources species could use per survey. We built presence absence models for all non-kleptoparasitic species that occurred sufficiently within our data set (the core community) using binomial GLMM’s with location as a random effect. We excluded kleptoparasites, as they do not depend on specific floral resources. For each species we only selected transect counts within their flight period, as retrieved from [30], to prevent including ‘meaningless’ 0’s. Model structure was similar to the models described above.

## 3. Results

### 3.1 Community composition

The median number of individuals caught per transects was 8, with totals varying between 0 and 384 (Appendix B.2). In total 12178 bees were recorded, of which 10503 were wild bees and 1675 honeybees. In total we found 156 species, with an estimated 199 occurring (Chao estimator, SE ± 21.68, see figure 3). Of these, the core community consisted of 39 species (89.4% total abundance) that occurred on 10% of all locations (Appendix C.2). This included five red-listed species (*Andrena gravida, A. labialis, A. wilkella, Nomada bifasciata and N. fulvicornis*), demonstrating that dikes can support vulnerable species. For 56 species, only 5 or fewer individuals were caught, with 25 species only caught once.

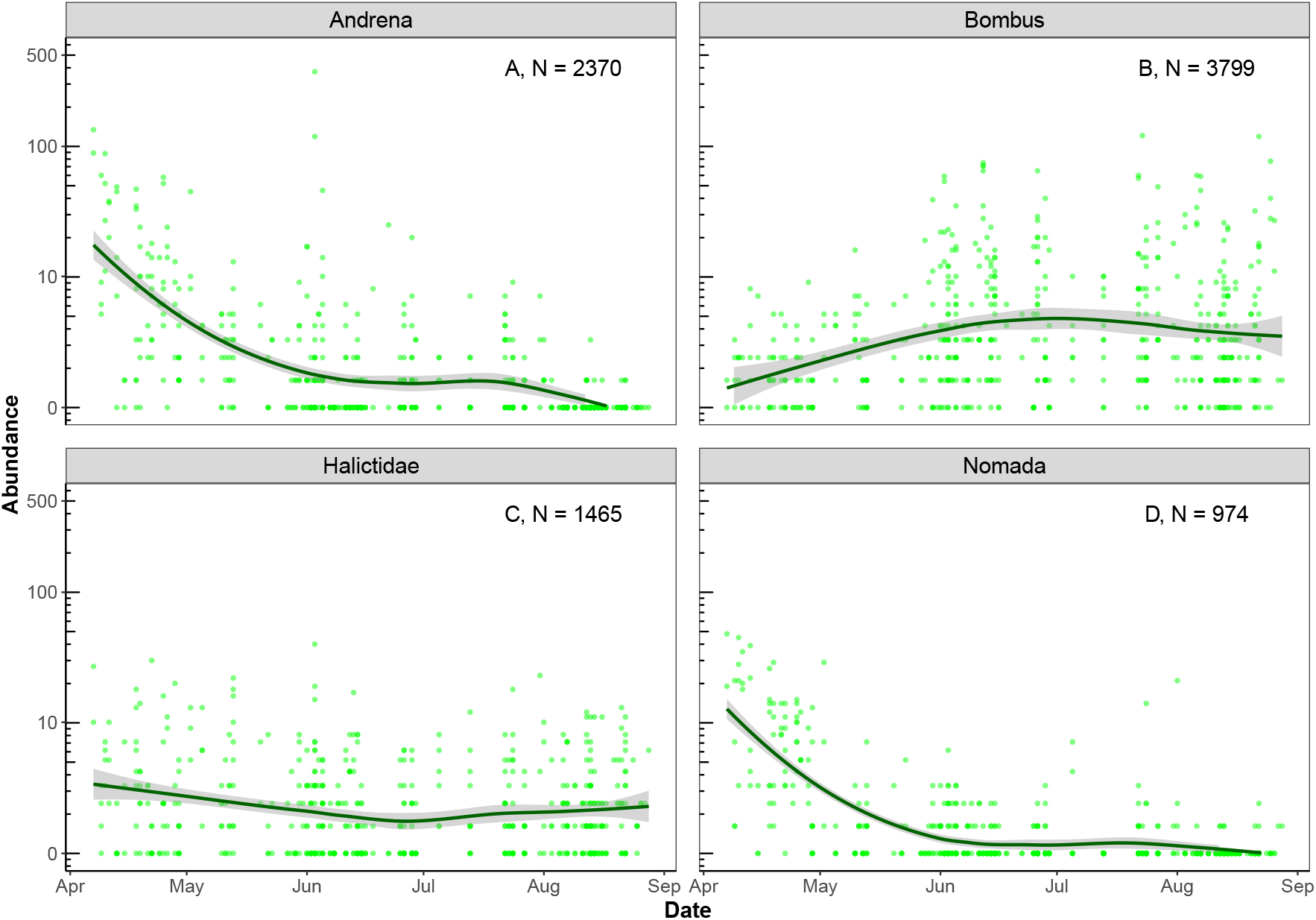

Most species belonged to the genera *Andrena* (2508 individuals), *Nomada* (1017 individuals), *Lasioglossum* + *Halictus* (1694 individuals, henceforth Halictidae) and *Bombus* (4534 individuals) (table 1, Appendix C.3 for all genera). The genus *Andrena* and Halictidae family consists of species that nest underground. *Nomada* and *Sphecodes* are kleptoparasitic bees, laying their eggs in nests of other bees and respectively having *Andrena* and Halictidae species as their main hosts. The omnipresence of kleptoparasites suggests that dikes are important nesting ground for wild bees. Concomitantly we only observed 25 above-ground nesting species (327 individuals), a low number compared to the soil nesting species (4441 individuals, *Bombus* excluded) representing percentage wise fewer species (table 1). Finally, 942 individuals (8.96% of the total) belonging to 37 species (23.72% of the total) are red-listed in the Netherlands (Appendix C.4).

**Table 1:** Most species-rich bee genera in the Netherlands, their total number of species occurring in the Netherlands and the number of species found in our surveys translated to the percentage of the total number of species. Their general nesting strategy is given in the fifth column.

| Taxon | #Species in the Netherlands | #Species on our dikes | Share of Dutch species found on dikes in this study (%) | Nesting |
| --- | --- | --- | --- | --- |
| *Andrena* | 75 | 37 | 49 | Below Ground |
| Halictidae | 52 | 22 | 42 | Below Ground |
| *Bombus* | 29 | 17 | 59 | Both |
| *Hoplites* + *Osmia* | 20 | 5 | 25 | Above Ground |
| *Megachile* | 13 | 5 | 38 | Above Ground |
| *Hylaeus* | 25 | 9 | 36 | Above Ground |
| *Nomada* | 48 | 22 | 46 | Parasitic |
| *Sphecodes* | 20 | 12 | 60 | Parasitic |
| Total Anthophila | 282 | 124 | 44 |  |

### 3.2 Phenology

Species activity was not distributed equally over the year (Figure 4, Appendix E.1 for other genera). The two most species-rich genera *Andrena* and *Nomada* were mostly active early during the year (Figure 4A and 4D), while *Bombus* is mostly active during summer (Figure 4B). The Halictidae group followed no clear pattern through time (Figure 4C). Similar activity patterns were observed for bee species richness (Appendix E.2). For red-listed species this pattern was even more pronounced, with only 59 of the total 942 observations after the 1^st^ of June (Appendix E.3).

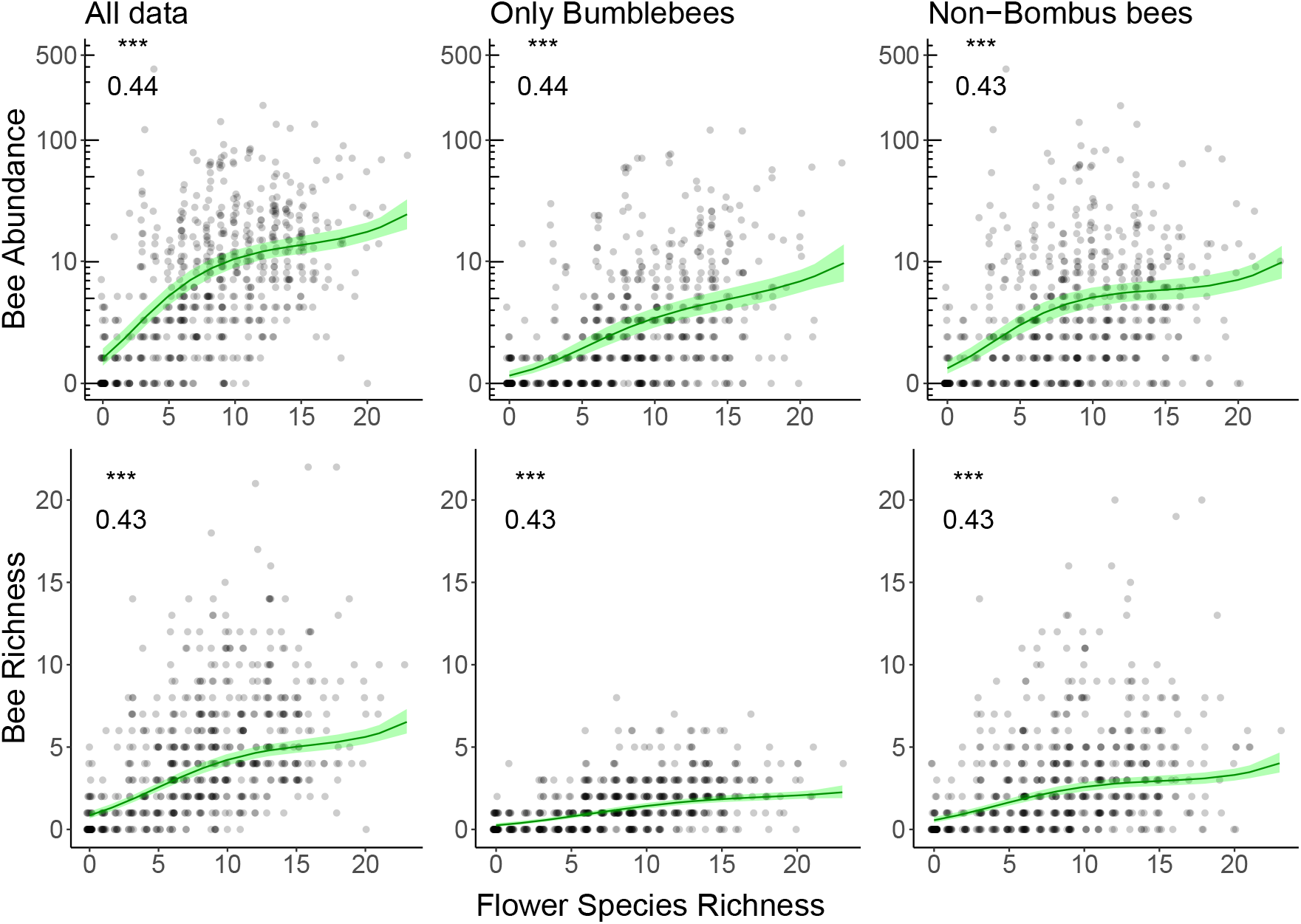

### 3.3 Relation Bee – Vegetation

Bee abundance and diversity in our dataset increased significantly with flower species richness in all considered subsets (figure 4, Appendix D.5 and D.6, marginal R^2^ presented in D.11 and D.12). This relationship is non-linear; third or fourth polynomial models describing flattening curves provided the best fit in all cases (figure 4, Appendix D.9 and D.10). Bee abundance and diversity increase rapidly at low flower species richness, after which curves flatten at around eight flower species. This is further supported by breakpoint analysis (Appendix A.4 and A.5).

The breakpoint was stronger for non-*Bombus* bees than bumblebees in our dataset, with slope for non-*Bombus* abundance and diversity changing from 0.15 and 0.33 before the breakpoint to 0.004 and 0.06 after respectively, compared to 0.14 and 0.16 to 0.09 and 0.05 respectively for bumblebees (figure 4, Appendix A.4 and A.5). This is consistent with the idea that generalist and highly mobile bumblebees keep increasing with floral resources, whereas other wild bees are limited by other factors after initial resource limitation is relieved.

Julian date showed a negative relationship with non-*Bombus* abundance and diversity, a positive effect on bumblebee abundance and no significant relationship with bumblebee diversity (Appendix D.7 and D.8), in line with the activity patterns described in 3.2. From the seven landscape factors, most parsimonious models for both abundance and diversity of all bees and non-*Bombus* bees included the area of NNN in a 1km radius (appendices D.9 and D.10). In these models a significant positive effects of more area NNN was found, although the model estimates were three to five times lower than the model estimated for local flower richness (Table 2). In addition, we observe a positive effect of area NNN in a 250m radius on total bee diversity. For bumblebees, model AIC’s were closer together and after correcting for the observer effect (Methods 2.3.3) we found no significant effects of landscape on bumblebee abundance (Table 2, Appendix D.13), and a small positive significant effect of area NNN in a 1km and 250m radius and a negative effect of increasing distance to these areas (Table 2).

**Table 2:** P-values (NS,*,**,***) and model estimates from models on different subsets of wild bees (in rows) for different landscape variables (columns). NNN stands for ‘NatuurNetwerk Nederland’, a network that includes green infrastructure and natural areas in the Netherlands. In grey the most parsimonious models (lowest AIC values). ^1^) Values for bumblebee models stem from GLM models correcting for observer (see methods 2.3.3). Grey squares indicate the model with the lowest AIC value for that model.

|  | NNN<br>in<br>1km | NNN<br>in<br>250m | Shortest<br>distance<br>to NNN | Agricultural<br>area in<br>250m | Agricultural<br>area in<br>1000m | Grassland<br>area in<br>250m | Grassland<br>area in<br>1000m |
| --- | --- | --- | --- | --- | --- | --- | --- |
| **All data**<br>**Abundance** | 0.15, * | 0.09,<br>NS | 0.06, NS | 0.09, NS | 0.12, NS | -0.02, NS | -0.04, NS |
| **Bumblebee**<br>**Abundance<sup>1</sup>** | 0.08,<br>NS | 0.09,<br>NS | -0.02,<br>NS | -0.05, NS | 0.11, NS | 0.002, NS | -0.08, NS |
| **Non-Bombus**<br>**bees**<br>**Abundance** | 0.20, * | 0.08,<br>NS | 0.09, NS | 0.12, NS | 0.14, NS | -0.08, NS | -0.03, NS |
| **All data**<br>**Diversity** | 0.11,<br>*** | 0.09,<br>** | -0.04,<br>NS | -0.05, NS | -0.03, NS | 0.02, NS | 0.03, NS |
| **Bumblebees**<br>**Diversity<sup>1</sup>** | 0.07, * | 0.08, * | -0.11, ** | -0.09, NS | -0.07, NS | 0.06, NS | 0.01, NS |
| **Non-Bombus**<br>**bees**<br>**Diversity** | 0.13,<br>** | 0.08,<br>NS | -0.001,<br>NS | -0.02, NS | -0.01, NS | -0.01, NS | 0.02, NS |
| **Red-listed**<br>**presence** | 0.41,<br>*** | 0.27, * | -0.09,<br>NS | -0.18, NS | -0.17, NS | 0.22, NS | 0.27, * |

Analysis of the presence of red-listed species shows that the chance of red listed individuals being present increases with increasing flower species richness (Appendix D.14 and D.15). Opposing overall patterns, we see that this chance increases without saturating effects. In addition, Julian date correlated negatively with red listed bee presence, suggesting that red-listed species in our system are primarily active in early spring. At the landscape level (Appendix 16 for AIC values), there was a strong positive effect of area NNN in a 1km radius, and smaller significant effect of grassland area in a 1km radius and area NNN in a 250m radius (Table 2).

### 3.4 Predicting species occurrence

For 20 of 25 species in the core community, we found a significant increase in their presence with increase of the number of their preferred flower species (figure 5, Appendix F.1). Five species form an exception to this. *Andrena vaga* had no food sources on dikes (therefore not included in figure 5), and four showed non-significant relationships (*Andrena gravida, A*. nitida, *Lasioglossum malachurum* and *L. xanthopus*). Another two had low marginal R-square values (*A. labialis* and *A. wilkella*), indicating a poor fit. These species are ground-nesting bees that we also observed nesting on our dikes.

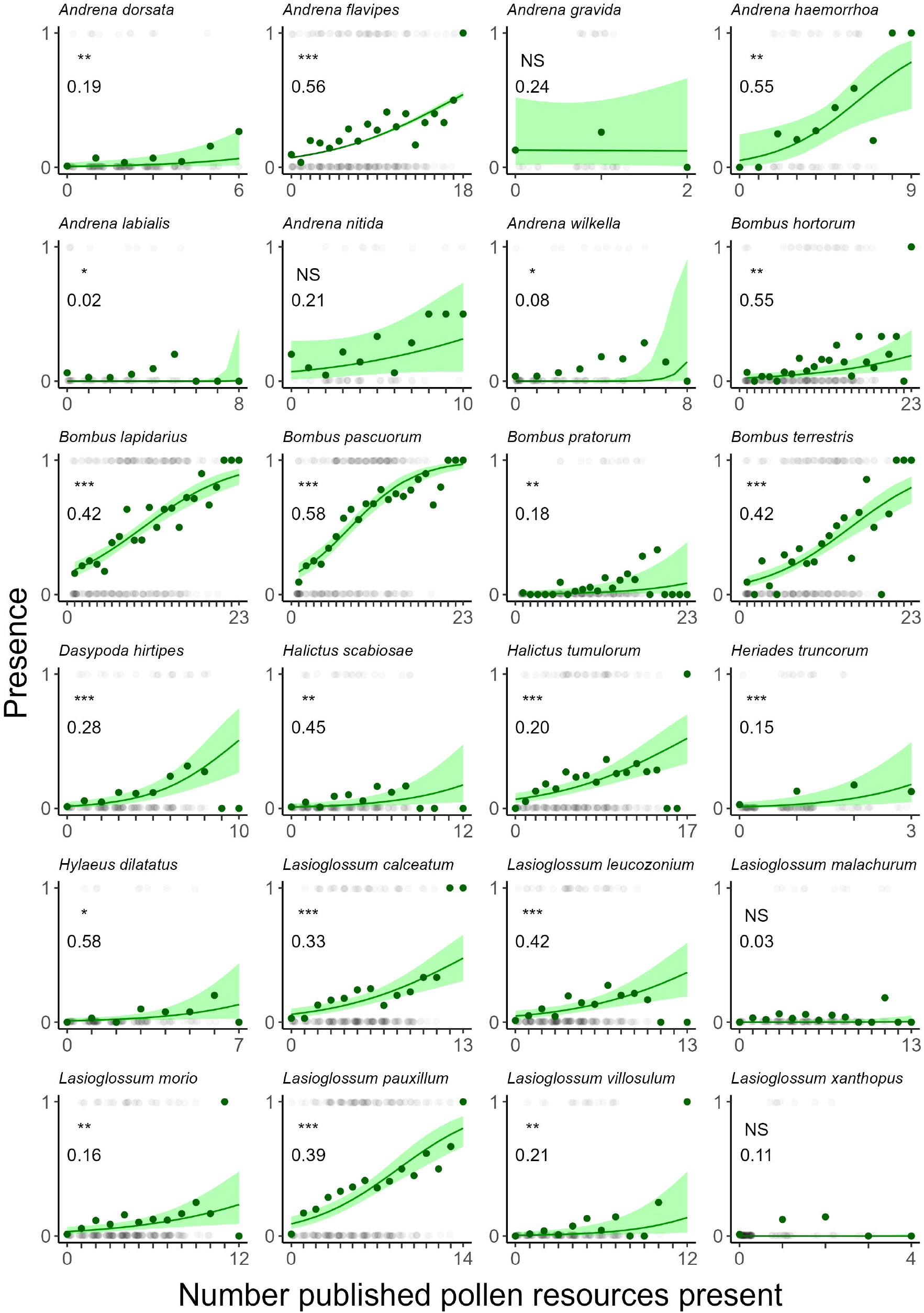

## 4. Discussion

Previous studies have shown that dikes can harbour rich plant, true bug and grasshopper communities [42–45]. By analysing an extensive dataset on bee occurrence on semi-natural dike grasslands in the Netherlands, we show that dikes support bee communities of large conservation interest. We encountered nearly half of all Dutch bee species, with nearly 10% of all individuals and over 20% of all species on the national red list. Local conditions on dike grasslands were the strongest predictor of bee diversity and abundance, underscoring that dike grasslands provide valuable bee habitat with both a nesting and resource function, even without the presence of natural areas nearby. Increase in flower-richness increased bee richness and abundance, but this effect saturated around eight flowering plant species present. This suggests that even modest increases in plant species richness can rapidly benefit wild bees. Nevertheless, the presence of red-listed species increased most after this saturation point, and the effect of natural area surrounding the location was stronger for these bees. This suggests that red-listed species rely on both landscape-level habitat presence and high local habitat quality, thus requiring conservation efforts that consider both landscape context and local needs of these rare species.

### 4.1 Benchmarking wild bee communities on dikes

Comparison with other studies from North-Western Europe on linear landscape elements further emphasises the value of dike grasslands. Firstly, an encountered 156 bee species (and an estimated 199) occurring on dikes seems high. Other inventories of bees in agricultural landscapes mostly report well below a 100 species even in countries that harbour more bee species in total, e.g. [46– 48]. In addition, we encountered high numbers of non-*Bombus* bees versus bumblebees (4969 vs. 4525 respectively) and red listed species (942 individuals). For instance, in flower strips in rural areas in the Netherlands, Scheper et al. (2015) found 5768 bumblebees compared to 1405 other wild bees, and only 13 individuals belonging to red listed species. Another study from Wood et al., (2015) found comparable species numbers and abundance proportions, but a lower number of individuals of conservation interest (114 individuals). Although we do not have a direct explanation, we do note that many of our grasslands are older compared to the systems referred to above, and meta-analysis has shown that pollinator community richness increases with restoration age [24]. In addition, the warm microclimate on sun-facing dike slopes is most likely favoured by bees. While comparisons between studies should be done with caution as sampling effort, sampling method and local context differ, dike grasslands do line up as habitat worth of special conservation interest for wild bees.

### 4.2 Local vegetation factors associated with species richness and abundance

Bee abundance and diversity in our dataset increased with flower species richness and abundance, a relationship described in previous studies [31]. This relationship was only apparent at low flower species richness, with bee abundance and diversity saturating around eight flower species. This saturating effect was much stronger for non-*Bombus* wild bees (comprising the majority of all bee diversity) compared to bumblebees. Whereas bumblebees are most represented in literature [50], this result shows that non-*Bombus* wild bees should be considered separately for effective conservation. This finding implies that conservation efforts aimed at increasing floral resources could quickly benefit wild bee communities as a whole on flower poor locations, although after being relieved of initial floral resource limitation other factors become limiting for non-*Bombus* wild bees.

Similar saturating relationships between flower species richness and pollinator diversity have been shown in the Jena experiment, where plant species richness is controlled in eighty 400m^2^ plots [51]. Although this saturating relationship could be expected for species diversity, we were surprised to also describe it for wild bee abundance. When increased flower species richness and abundance does not lead to an increase in pollinator abundance, this will essentially lead to fewer pollinating individuals per flowering plant. This relationship has been described by Fantinato et al., 2021, who observed a humped-shaped response of pollinator visitation to floral resource availability. These results are consistent with a recent study linking declines in insect-pollinated plant species in the Netherlands to widespread pollinator decline [53]. Follow-up research on the effects of pollinator saturation on pollination success of flowers would be valuable evidence for this idea.

In contrast to the overall saturating relationships, the occurrence of red-listed species in our dataset increased most at higher flower richness. Red-listed species are known to show a more selective response to habitat characteristics [54]. An increase in flower species richness could hence increase the chance of host flower presence and subsequently red-listed bee species occurrence. Nevertheless, specialist bees in our dataset mostly rely on more common flower species, a trait commonly reported as nested interaction patterns in scientific literature [55]. An additional mechanism through which increased flower species richness could promote red-listed bee species occurrence can be found in niche partitioning. Computational models suggest that an increase in flower diversity allows bees to partition their resource niches, benefiting species with weaker competitive strength and/or more specific resource preferences as generalist species shift their collection patterns to other flower species [56]. Although study of the exact mechanism behind this requires further analysis of interaction patterns at our dikes, our results suggest that red-listed species conservation requires a different approach from the general bee community.

Further analysis on the occurrence of individual species revealed generally strong relationships with the occurrence of their known pollen host flowers. Hence, not only flower species richness, but also the identity of the flowering species determines the success of conservation efforts [57–60]. Our results thus open the possibility that in addition to general community richness, conservation effort may steer towards the conservation of specific bee species through measures aimed at promoting specific host flower species.

### 4.3 Landscape setting

Previous studies have shown the importance of landscape setting for local bee community composition (e.g. Griffin et al., 2021; Scheper et al., 2015). Wild bees use partial habitats, with nesting locations being separate from foraging locations. In addition, more natural area can function as reservoirs from which species can find their way to the studied location [63] or as buffer when local conditions are unfavourable (e.g. after mowing). While our data did not allow investigating how detailed landscape structure (e.g. availability of above-ground nesting places) was related to local bee communities, analysis of separate landscape elements revealed a small but consistent positive effect of more landscape level natural area, primarily on non-*Bombus* bees. In our dataset we find a number of above-ground breeding species (e.g. *Chelostoma florisomne* and *Hylaeus dilatatus*) that must depend on surrounding areas as there is limited to no suitable vegetation growing on dikes. In addition, for red-listed species the positive effect of natural area was only slightly smaller than that of flower species richness. However, at the community level the positive effect of natural area in the surroundings was three to five times smaller than the effect of local flower richness. In addition, the occurrence of most of the common species could accurately be predicted through the presence of their preferred resources. This suggests that dike grasslands can sustain bee communities even in otherwise heavily altered landscapes. We thus conclude that local conservation efforts on dikes are worthwhile regardless of landscape context.*4*.*4 Temporal differences between solitary bees and bumblebees*

Activity of most wild bee species on our grasslands, and especially both species richness and abundance of red-listed bee species, was skewed towards early spring, notably dropping after mid-June (figure 3). These results indicate that dike grasslands are specifically valuable early spring habitats. This is contradicting the pattern that most bee species in the Netherlands are active during June through August [64]. We hypothesize that the warm sun facing dike slopes are likely ideal nesting locations especially in early spring when average temperatures are still low. In addition, local bee communities on dikes might adapt to long term stable management of dike grasslands, where mowing typically happens in June.

Compared to the higher abundance and diversity of non-*Bombus* bees in spring, we observed inversed temporal trends for bumblebees. Bumblebee colonies usually grow during the year and as such higher numbers can be expected in summer. We see two possible explanations for the lower number of non-*Bombus* bees later in the year. Firstly, this could be an example of temporal niche differentiation, where species avoid each other in time to reduce competition [65]. Literature does indeed suggest that competition for floral resources plays an important role in bee community formation, making this explanation plausible [66]. However, it could also be the outcome of long-term mowing management at most dikes in our dataset. Bumblebees are less tied to the dike specifically, due to their long flight ranges. As such, mowing regime will have a limited effect on their colonies. Non-*Bombus* bees are often tied to specific locations, making it harder to deal with mowing events.

### 4.5 Dike grasslands as nesting habitat for bees

Next to floral resources, bees require a place to make their nest. In this study, several observations suggest that dike grasslands also function as nesting location for ground-nesting bee species. Firstly, we observed especially large numbers of ground-nesting bees and their associated kleptoparasites, while aboveground nesting species were remarkably scarce. The high abundance and diversity of kleptoparasites would likely not occur without stable nesting communities of their host species. In addition, we find several diverse and abundant bee communities at low flower richness, where we also observed bee nesting colonies in the field. Finally, although most species show strong relationships with their host plant species presence in the dike grassland, several ground nesting bee species did not. *Andrena vaga* is a clear example, being found at more than 10% of all locations, suggesting that the species has its nesting location on dikes and its food source (*Salix* spp, willows) nearby. Their role as valued nesting habitat could be an additional reason as to why dikes harbour such rich wild bee communities.

If the availability of nesting habitat is important to consider in conservation projects [49,67], this raises the question how increases in flower species richness and vegetation structure (including open spots for nesting) are connected. Higher plant diversity has been linked to more vegetation structure complexity, but the general pattern is suggested to be saturating at modest plant diversity [68]. At a certain point increased plant diversity most likely comes with increased vegetation coverage, which has been shown to have a negative relationship with bee nesting in artificially created bare soil plots [69]. To our knowledge, direct studies on the relationship between VSC and bee nesting do not exist, making it difficult to speculate whether the saturating patters in non-*Bombus* diversity in our study can be explained through saturation in VSC and thus nesting availability. In any way we suggest that how nesting and resource functions can best be combined in conservation efforts is an important topic for future studies.

## 5. Conclusions

Linear landscape elements are known as valuable area for pollinator conservation [70,71]. Our analysis highlights the unique potential of dike grasslands for wild bee conservation, with rich early-spring bee communities and many red-listed species that distinguishes them from many other linear landscape elements. The saturating patterns that we report indicate that, at poor dikes, modest conservation efforts can quickly benefit wild bee communities. In addition, the strong effect of local vegetation suggests that conservation efforts, regardless of landscape context, will have positive effects. Future research should aim to study how nesting and resource functions can best be combined.

## 6. Data Statement

Research data will be archived open-access in a Data Repository like Dryad, Zenodo or DANS. To aid reviewers, we have added our data file to the submitted manuscript files.

## 7. Conflict of interests

Authors confirm that they had no conflict of interests in this study. This research did not receive any specific grant from funding agencies in the public, commercial, or not-for-profit sectors.

## 8. Acknowledgements

The authors are grateful to all students and co-workers that helped to collect the data throughout this project: Ivanka Spruijt, Lisa van den Dries, Maarten van Emmerik, Lars Willighagen and Else Dieks (all Radboud University), Miriam Scherpenisse, Jochem Macaré and Jeroen Willems (Natuurbalans) and Cyril Liebrand (Eureco). In addition, the authors extend their gratitude to the waterboards Rivierenland, Aa en Maas, Waterschap Hollandse Delta and Waterschap Brabantse Delta as well as dike managers Natuurmonumenten, Staatsbosbeheer, Het Zuid-Hollands Landschap, Het Zeeuwse Landschap, de Ploegdriever and Marc Büchner for allowing the collection of the data on their dike grasslands.

## 9. Author contributions

CS: Conceptualization, Methodology, Investigation, Formal Analysis, Writing – original draft. LS, IR, NG: Methodology, Investigation, Writing – Review & Editing. EJ: Formal Analysis, Writing – Review & Editing. HdK: Supervision, Writing – Review & Editing.

## Notes

### Competing Interest Statement

The authors have declared no competing interest.

