## Appendix C for "Phenology and flower species availability define wild bee communities on river embankments"

**Appendix C – Community description**

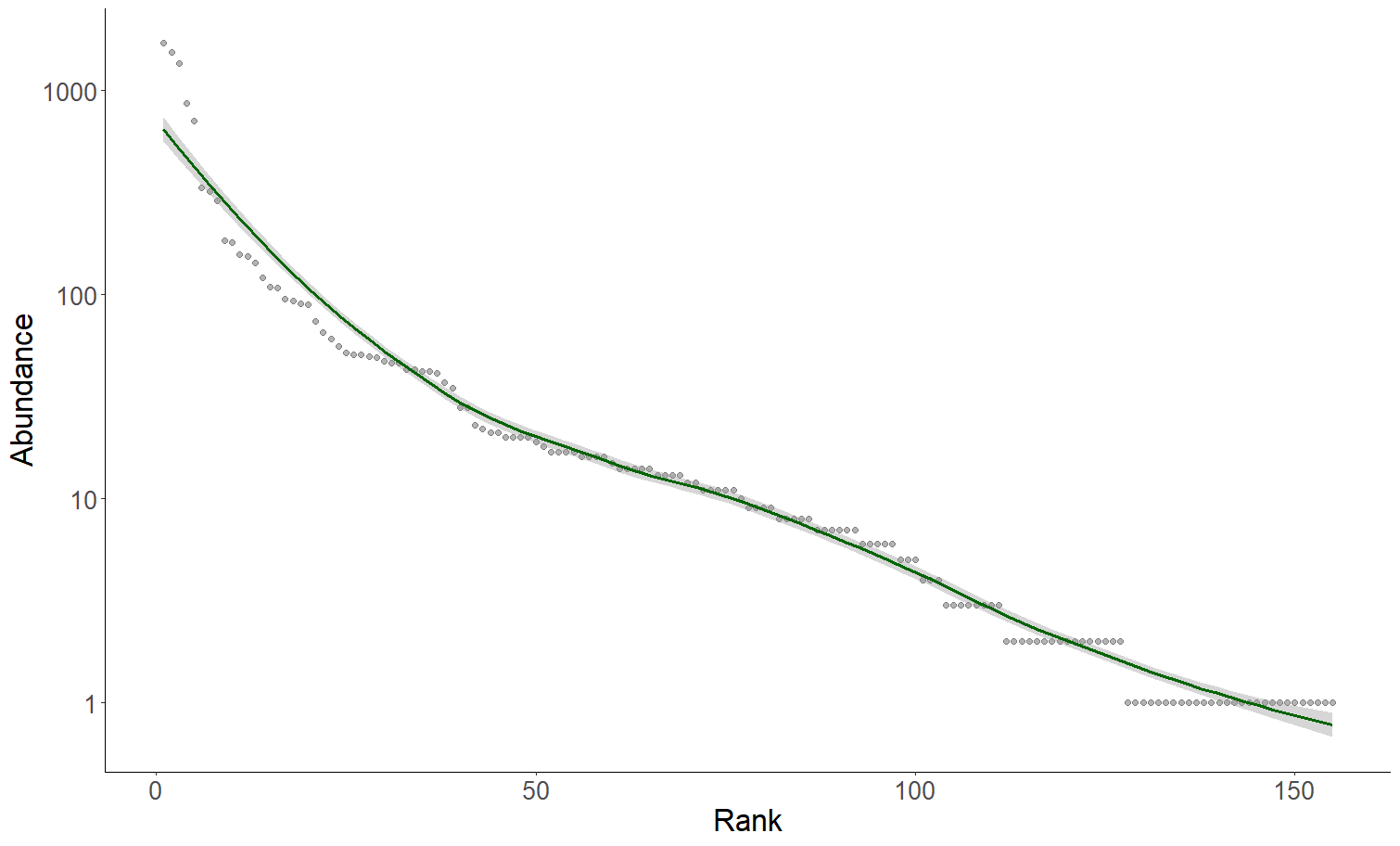

Appendix C.1: Rank abundance curve of all species in our data set on a log scale. The dark green loess curve was added to aid visual interpretation.

Appendix C.2*:* list of species that occur at more than 10% of all locations, with the number of locations they occur at and their abundance (Quantity). Species in bold are red listed.

| ***Species*** | ***Number of locations*** | ***Quantity*** | ***Species*** | ***Number of locations*** | ***Quantity*** |
| --- | --- | --- | --- | --- | --- |
| *Bombus lapidarius* | *130* | *1718* | *Nomada marshamella* | *27* | *46* |
| *Bombus pascuorum* | *124* | *1554* | *Sphecodes monilicornis* | *27* | *56* |
| *Bombus terrestris* | *105* | *872* | *Andrena dorsata* | *26* | *74* |
| *Lasioglossum pauxillum* | *102* | *715* | ***Nomada bifasciata*** | *25* | *154* |
| *Andrena flavipes* | *100* | *1355* | *Nomada fabriciana* | *25* | *49* |
| *Lasioglossum calceatum* | *74* | *185* | *Andrena nitida* | *23* | *50* |
| *Halictus tumulorum* | *73* | *289* | *Lasioglossum villosulum* | *23* | *53* |
| *Nomada fucata* | *67* | *333* | *Bombus pratorum* | *23* | *46* |
| *Lasioglossum leucozonium* | *52* | *111* | ***Nomada fulvicornis*** | *22* | *51* |
| *Lasioglossum morio* | *51* | *93* | ***Andrena wilkella*** | *20* | *51* |
| *Bombus hortorum* | *46* | *158* | *Nomada goodeniana* | *20* | *28* |
| *Dasypoda hirtipes* | *41* | *181* | *Bombus campestris* | *19* | *43* |
| *Andrena haemorrhoa* | *40* | *143* | *Lasioglossum malachurum* | *17* | *42* |
| *Halictus scabiosae* | *30* | *109* | *Bombus hypnorum* | *16* | *47* |
| *Nomada ruficornis* | *30* | *42* | *Lasioglossum xanthopus* | *16* | *37* |
| ***Andrena gravida*** | *28* | *321* | *Nomada lathburiana* | *16* | *65* |
| *Heriades truncorum* | *28* | *90* | ***Andrena labialis*** | *15* | *41* |
| *Nomada flava* | *28* | *89* | *Andrena minutula* | *15* | *28* |
| *Andrena vaga* | *27* | *95* | *Hylaeus dilatatus* | *15* | *20* |
| *Nomada flavoguttata* | *27* | *61* |  |  |  |

Appendix C.3: Number of species per genus

| **Genus** | **Quantity** | **Genus** | **Quantity** |
| --- | --- | --- | --- |
| Andrena | 37 | Melitta | 3 |
| Nomada | 22 | Osmia | 3 |
| Lasioglossum | 18 | Anthidiellum | 1 |
| Bombus | 17 | Anthophora | 1 |
| Sphecodes | 12 | Ceratina | 1 |
| Hylaeus | 9 | Chalicodoma | 1 |
| Colletes | 5 | Coelioxys | 1 |
| Halictus | 5 | Dasypoda | 1 |
| Megachile | 5 | Epeoloides | 1 |
| Stelis | 4 | Epeolus | 1 |
| Chelostoma | 3 | Heriades | 1 |
| Hoplitis | 3 | Macropis | 1 |

Appendix C.4: Red listed species and their abundance in our dataset

| **Species** | **Quantity** | **Species** | **Quantity** |
| --- | --- | --- | --- |
| Andrena gravida | 321 | Bombus jonellus | 3 |
| Nomada bifasciata | 154 | Lasioglossum lativentre | 3 |
| Andrena bimaculata | 121 | Stelis ornatula | 3 |
| Andrena wilkella | 52 | Andrena trimmerana | 2 |
| Nomada fulvicornis | 51 | Bombus sylvarum | 2 |
| Andrena labialis | 41 | Hoplitis tridentata | 2 |
| Bombus vestalis | 35 | Nomada integra | 2 |
| Andrena rosae | 17 | Andrena fulvago | 1 |
| Bombus ruderarius | 17 | Andrena semilaevis | 1 |
| Nomada stigma | 17 | Andrena varians | 1 |
| Osmia niveata | 16 | Bombus barbutellus | 1 |
| Andrena hattorfiana | 15 | Bombus veteranus | 1 |
| Andrena humilis | 11 | Hoplitis claviventris | 1 |
| Sphecodes spinulosus | 11 | Hylaeus pectoralis | 1 |
| Halictus quadricinctus | 9 | Megachile lapponica | 1 |
| Andrena ovatula | 8 | Nomada striata | 1 |
| Bombus rupestris | 8 | Stelis breviuscula | 1 |
| Bombus muscorum | 7 | Stelis phaeoptera | 1 |
| Andrena lathyri | 3 |  |  |
