## Appendix D for "Phenology and flower species availability define wild bee communities on river embankments"

**Appendix D - Models**

Appendix D.1; Significance outcomes (NS, *, **, ***) of Kruskal-Wallis tests on the residuals of each wild bee abundance model grouped by observer. Each cell represents a model, with the dataset used in the first column and the polynomial in the second. Other columns indicate the landscape factor incorporated in the model.

|  | **Poly-nomial** | **NNN in 1km** | **NNN in 250m** | **Shortest distance to NNN** | **Agri area in 250m** | **Agri area in 1000m** | **Grass area in 250m** | **Grassland area in 1000m** | **No landscape factor** |
| --- | --- | --- | --- | --- | --- | --- | --- | --- | --- |
| **All data** | 1 | * | * | ** | * | * | * | * | ** |
|  | 2 | * | NS | * | * | * | * | * | * |
|  | 3 | NS | NS | NS | NS | NS | NS | NS | NS |
|  | 4 | NS | NS | NS | NS | NS | NS | NS | NS |
| **Only bumble-bees** | 1 | ** | ** | ** | ** | ** | ** | ** | *** |
|  | 2 | ** | ** | ** | ** | ** | ** | ** | ** |
|  | 3 | ** | ** | ** | ** | ** | ** | ** | ** |
|  | 4 | ** | ** | ** | ** | ** | ** | ** | ** |
| **Non-Bombus bees** | 1 | NS | NS | NS | NS | NS | NS | NS | NS |
|  | 2 | NS | NS | NS | NS | NS | NS | NS | NS |
|  | 3 | NS | NS | NS | NS | NS | NS | NS | NS |
|  | 4 | NS | NS | NS | NS | NS | NS | NS | NS |

Appendix D.2; Significance outcomes (NS, *, **, ***) of Kruskal-Wallis tests on the residuals of each wild bee diversity model grouped by observer. Each cell represents a model, with the dataset used in the first column and the polynomial in the second. Other columns indicate the landscape factor incorporated in the model.

|  | **Poly-nomial** | **NNN in 1km** | **NNN in 250m** | **Shortest distance to NNN** | **Agri area in 250m** | **Agri area in 1000m** | **Grass area in 250m** | **Grassland area in 1000m** | **No landscape factor** |
| --- | --- | --- | --- | --- | --- | --- | --- | --- | --- |
| **All data** | 1 | NS | NS | NS | NS | NS | NS | NS | NS |
|  | 2 | NS | NS | NS | NS | NS | NS | NS | NS |
|  | 3 | NS | NS | NS | NS | NS | NS | NS | NS |
|  | 4 | NS | NS | NS | NS | NS | NS | NS | NS |
| **Only bumble-bees** | 1 | *** | *** | *** | ** | ** | *** | *** | *** |
|  | 2 | *** | *** | *** | ** | ** | ** | *** | *** |
|  | 3 | *** | ** | *** | ** | ** | ** | *** | *** |
|  | 4 | *** | ** | *** | ** | ** | ** | *** | *** |
| **Non-Bombus bees** | 1 | NS | NS | NS | NS | NS | NS | NS | NS |
|  | 2 | NS | NS | NS | NS | NS | NS | NS | NS |
|  | 3 | NS | NS | NS | NS | NS | NS | NS | NS |
|  | 4 | NS | NS | NS | NS | NS | NS | NS | NS |

Appendix D.3: P-values (NS,*,**,***) and model estimates for different landscape variables in models on bee abundance from three datasets (in rows) incorporating different landscape level factors (columns). NNN stands for ‘NatuurNetwerk Nederland’, a network that includes green infrastructure and natural areas in the Netherlands. In grey the most parsimonious models (lowest AIC values).

|  | **NNN in 1km** | **NNN in 250m** | **Shortest distance to NNN** | **Agricultural area in 250m** | **Agricultural area in 1000m** | **Grassland area in 250m** | **Grassland area in 1000m** |
| --- | --- | --- | --- | --- | --- | --- | --- |
| **All data** | 0.15, * | 0.09, NS | 0.06, NS | 0.09, NS | 0.12, NS | -0.02, NS | -0.04, NS |
| **Only bumblebees** | 0.18, * | 0.21, * | -0.14, NS | -0.15, NS | -0.13, NS | 0.19, * | 0.05, NS |
| **Non-Bombus bees** | 0.20, * | 0.08, NS | 0.09, NS | 0.12, NS | 0.14, NS | -0.08, NS | -0.03, NS |

Appendix D.4: P-values (NS,*,**,***) and model estimates for different landscape variables in models on bee diversity from three datasets (in rows) incorporating different landscape level factors (columns). NNN stands for ‘NatuurNetwerk Nederland’, a network that includes green infrastructure and natural areas in the Netherlands. In grey the most parsimonious models (lowest AIC values).

|  | **NNN in 1km** | **NNN in 250m** | **Shortest distance to NNN** | **Agricultural area in 250m** | **Agricultural area in 1000m** | **Grassland area in 250m** | **Grassland area in 1000m** |
| --- | --- | --- | --- | --- | --- | --- | --- |
| **All data** | 0.11, *** | 0.09, ** | -0.04, NS | -0.05, NS | -0.03, NS | 0.02, NS | 0.03, NS |
| **Only bumblebees** | 0.11, ** | 0.13, ** | -0.14, ** | -0.14, *** | -0.14, *** | 0.11, ** | 0.07, NS |
| **Non-Bombus bees** | 0.13, ** | 0.08, NS | -0.001, NS | -0.02, NS | -0.01, NS | -0.01, NS | 0.02, NS |

Appendix D.5: P-values (NS,*,**,***) and model estimates for flower species richness in models on bee abundance from three datasets (in rows) incorporating different landscape level factors (columns). NNN stands for ‘NatuurNetwerk Nederland’, a network that includes green infrastructure and natural areas in the Netherlands. In grey the most parsimonious models (lowest AIC values).

|  | **NNN in 1km** | **NNN in 250m** | **Shortest distance to NNN** | **Agricultural area in 250m** | **Agricultural area in 1000m** | **Grassland area in 250m** | **Grassland area in 1000m** |
| --- | --- | --- | --- | --- | --- | --- | --- |
| **All data** | 0.84, *** | 0.85, *** | 0.86, *** | 0.86, *** | 0.86, *** | 0.86, *** | 0.86, *** |
| **Only bumblebees** | 0.96, *** | 0.96, *** | 0.97, *** | 0.97, *** | 0.97, *** | 0.97, *** | 0.97, *** |
| **Non-Bombus bees** | 0.70, *** | 0.71, *** | 0.72, *** | 0.73, *** | 0.73, *** | 0.72, *** | 0.72, *** |

Appendix D.6: P-values (NS,*,**,***) and model estimates for flower species richness in models on bee diversity from three datasets (in rows) incorporating different landscape level factors (columns). NNN stands for ‘NatuurNetwerk Nederland’, a network that includes green infrastructure and natural areas in the Netherlands. In grey the most parsimonious models (lowest AIC values).

|  | **NNN in 1km** | **NNN in 250m** | **Shortest distance to NNN** | **Agricultural area in 250m** | **Agricultural area in 1000m** | **Grassland area in 250m** | **Grassland area in 1000m** |
| --- | --- | --- | --- | --- | --- | --- | --- |
| **All data** | 0.54, *** | 0.54, *** | 0.55, *** | 0.55, *** | 0.55, *** | 0.55, *** | 0.55, *** |
| **Only bumblebees** | 0.62, *** | 0.62, *** | 0.63, *** | 0.62, *** | 0.62, *** | 0.63, *** | 0.63, *** |
| **Non-Bombus bees** | 0.50, *** | 0.51, *** | 0.52, *** | 0.51, *** | 0.52, *** | 0.52, *** | 0.52, *** |

Appendix D.7: P-values (NS,*,**,***) and model estimates for Julian date in models on bee abundance from three datasets (in rows) incorporating different landscape level factors (columns). NNN stands for ‘NatuurNetwerk Nederland’, a network that includes green infrastructure and natural areas in the Netherlands. In grey the most parsimonious models (lowest AIC values).

|  | **NNN in 1km** | **NNN in 250m** | **Shortest distance to NNN** | **Agricultural area in 250m** | **Agricultural area in 1000m** | **Grassland area in 250m** | **Grassland area in 1000m** |
| --- | --- | --- | --- | --- | --- | --- | --- |
| **All data** | -0.30, *** | -0.31, *** | -0.31, *** | -0.31, *** | -0.31, *** | -0.31, *** | -0.31, *** |
| **Only bumblebees** | 0.37, *** | 0.37, *** | 0.35, *** | 0.35, *** | 0.35, *** | 0.35, *** | 0.35, *** |
| **Non-Bombus bees** | -0.62, *** | -0.63, *** | -0.63, *** | -0.62, *** | -0.62, *** | -0.62, *** | -0.63, *** |

Appendix D.8: P-values (NS,*,**,***) and model estimates for Julian date in models on bee diversity from three datasets (in rows) incorporating different landscape level factors (columns). NNN stands for ‘NatuurNetwerk Nederland’, a network that includes green infrastructure and natural areas in the Netherlands. In grey the most parsimonious models (lowest AIC values).

|  | **NNN in 1km** | **NNN in 250m** | **Shortest distance to NNN** | **Agricultural area in 250m** | **Agricultural area in 1000m** | **Grassland area in 250m** | **Grassland area in 1000m** |
| --- | --- | --- | --- | --- | --- | --- | --- |
| **All data** | -0.30, *** | -0.30, *** | -0.31, *** | -0.31, *** | -0.31, *** | -0.31, *** | -0.31, *** |
| **Only bumblebees** | 0.04, NS | 0.03, NS | 0.03, NS | 0.01, NS | 0.01, NS | 0.02, NS | 0.02, NS |
| **Non-Bombus bees** | -0.46, *** | -0.46, *** | -0.47, *** | -0.47, *** | -0.47, *** | -0.46, *** | -0.47, *** |

Appendix D.9: AIC values of models on bee diversity from different datasets and polynomials (in rows) incorporating different landscape level factors or no landscape factor (columns). NNN stands for ‘NatuurNetwerk Nederland’, a network that includes green infrastructure and natural areas in the Netherlands. In grey the most parsimonious models (lowest AIC values).

|  | **Poly-nomial** | **NNN in 1km** | **NNN in 250m** | **Shortest distance to NNN** | **Agri in 250m** | **Agri in 1km** | **Grass in 250m** | **Grass in 1km** | **No factor** |
| --- | --- | --- | --- | --- | --- | --- | --- | --- | --- |
| **All data** | 1 | 2856.01 | 2861.14 | 2867.52 | 2867.95 | 2868.55 | 2868.78 | 2868.58 | 2866.98 |
|  | 2 | 2798.47 | 2802.44 | 2807.13 | 2807.25 | 2808.44 | 2808.83 | 2808.76 | 2807.07 |
|  | 3 | 2780.10 | 2785.45 | 2791.28 | 2790.88 | 2792.12 | 2792.77 | 2792.51 | 2791.13 |
|  | 4 | 2778.12 | 2783.23 | 2789.86 | 2789.11 | 2790.42 | 2791.30 | 2790.97 | 2789.85 |
| **Only bumble-bees** | 1 | 1847.23 | 1845.03 | 1845.94 | 1845.51 | 1845.69 | 1847.72 | 1851.92 | 1852.39 |
|  | 2 | 1814.25 | 1811.32 | 1810.98 | 1808.39 | 1810.07 | 1812.59 | 1817.71 | 1818.46 |
|  | 3 | 1811.80 | 1809.24 | 1809.42 | 1806.40 | 1807.93 | 1810.80 | 1815.83 | 1816.86 |
|  | 4 | 1813.66 | 1811.20 | 1811.33 | 1808.37 | 1809.91 | 1812.73 | 1817.74 | 1818.69 |
| **Non-Bombus bees** | 1 | 2515.40 | 2520.62 | 2523.79 | 2523.78 | 2523.71 | 2523.66 | 2523.73 | 2521.79 |
|  | 2 | 2489.72 | 2494.11 | 2496.62 | 2496.54 | 2496.59 | 2496.49 | 2496.60 | 2494.63 |
|  | 3 | 2475.28 | 2480.84 | 2483.96 | 2483.78 | 2483.96 | 2483.89 | 2483.81 | 2481.96 |
|  | 4 | 2471.54 | 2477.10 | 2480.97 | 2480.57 | 2480.92 | 2480.96 | 2480.65 | 2578.97 |

Appendix D.10: AIC values of models on bee abundance from different datasets and polynomials (in rows) incorporating different landscape level factors or no landscape factor (columns). NNN stands for ‘NatuurNetwerk Nederland’, a network that includes green infrastructure and natural areas in the Netherlands. In grey the most parsimonious models (lowest AIC values).

|  | **Poly-nomial** | **NNN in 1km** | **NNN in 250m** | **Shortest distance to NNN** | **Agri area in 250m** | **Agri area in 1000m** | **Grass area in 250m** | **Grassland area in 1000m** | **No landscape factor** |
| --- | --- | --- | --- | --- | --- | --- | --- | --- | --- |
| **All data** | 1 | 4191.96 | 4194.89 | 4195.42 | 4193.26 | 4191.33 | 4196.02 | 4195.72 | 4194.43 |
|  | 2 | 4164.48 | 4166.78 | 4166.93 | 4165.61 | 4163.74 | 4167.61 | 4167.09 | 4165.95 |
|  | 3 | 4152.63 | 4155.85 | 4156.59 | 4155.61 | 4154.13 | 4157.39 | 4157.10 | 4155.53 |
|  | 4 | 4152.77 | 4156.10 | 4157.34 | 4156.52 | 4155.13 | 4158.09 | 4157.85 | 4156.16 |
| **Only bumble-bees** | 1 | 3051.75 | 3050.44 | 3053.19 | 3053.60 | 3054.24 | 3051.25 | 3055.83 | 3053.91 |
|  | 2 | 3040.51 | 3038.94 | 3041.40 | 3041.08 | 3041.99 | 3039.11 | 3044.19 | 3042.32 |
|  | 3 | 3036.78 | 3035.63 | 3038.57 | 3037.88 | 3038.73 | 3035.62 | 3041.13 | 3039.45 |
|  | 4 | 3038.58 | 3037.31 | 3040.40 | 3039.63 | 3040.50 | 3037.36 | 3042.98 | 3041.34 |
| **Non-Bombus bees** | 1 | 3367.46 | 3372.08 | 3371.43 | 3369.56 | 3368.68 | 3371.60 | 3372.97 | 3371.04 |
|  | 2 | 3351.88 | 3355.87 | 3354.90 | 3353.41 | 3352.39 | 3355.12 | 3356.39 | 3354.53 |
|  | 3 | 3341.79 | 3346.77 | 3346.40 | 3345.66 | 3344.90 | 3346.99 | 3347.88 | 3345.89 |
|  | 4 | 3342.14 | 3347.36 | 3347.40 | 3346.80 | 3346.10 | 3348.03 | 3348.75 | 3346.75 |

Appendix D.11: Marginal R^2^ values for models on bee abundance from three datasets (in rows) incorporating different landscape level factors (columns). NNN stands for ‘NatuurNetwerk Nederland’, a network that includes green infrastructure and natural areas in the Netherlands. In grey the most parsimonious models (lowest AIC values).

|  | **NNN in 1km** | **NNN in 250m** | **Shortest distance to NNN** | **Agricultural area in 250m** | **Agricultural area in 1000m** | **Grassland area in 250m** | **Grassland area in 1000m** |
| --- | --- | --- | --- | --- | --- | --- | --- |
| **All data** | 0.44 | 0.43 | 0.43 | 0.43 | 0.44 | 0.43 | 0.43 |
| **Only bumblebees** | 0.44 | 0.44 | 0.42 | 0.43 | 0.43 | 0.43 | 0.42 |
| **Non-Bombus bees** | 0.43 | 0.42 | 0.42 | 0.42 | 0.42 | 0.42 | 0.41 |

Appendix D.12: Marginal R^2^ values for models on bee diversity from three datasets (in rows) incorporating different landscape level factors (columns). NNN stands for ‘NatuurNetwerk Nederland’, a network that includes green infrastructure and natural areas in the Netherlands. In grey the most parsimonious models (lowest AIC values).

|  | **NNN in 1km** | **NNN in 250m** | **Shortest distance to NNN** | **Agricultural area in 250m** | **Agricultural area in 1000m** | **Grassland area in 250m** | **Grassland area in 1000m** |
| --- | --- | --- | --- | --- | --- | --- | --- |
| **All data** | 0.33 | 0.33 | 0.32 | 0.32 | 0.32 | 0.32 | 0.32 |
| **Only bumblebees** | 0.32 | 0.32 | 0.32 | 0.32 | 0.32 | 0.31 | 0.31 |
| **Non-Bombus bees** | 0.36 | 0.35 | 0.34 | 0.34 | 0.34 | 0.34 | 0.34 |

Appendix D.13: P-values (NS,*,**,***) and model estimates for different landscape variables in models on bee abundance from three datasets (in rows) incorporating different landscape level factors (columns). NNN stands for ‘NatuurNetwerk Nederland’, a network that includes green infrastructure and natural areas in the Netherlands. In grey the most parsimonious models (lowest AIC values).

| **Landscape  factor**  **Model** | **NNN in 1km** | **NNN in 250m** | **Shortest distance to NNN** | **Agricultural area in 250m** | **Agricultural area in 1000m** | **Grassland area in 250m** | **Grassland area in 1000m** |
| --- | --- | --- | --- | --- | --- | --- | --- |
| **GLM Bumblebee Abudance** | 0.08, NS | 0.09, NS | -0.02, NS | -0.05, NS | 0.11, NS | 0.002, NS | -0.08, NS |
| **GLM Bumblebees Diversity** | 0.07, * | 0.08, * | -0.11, ** | -0.09, NS | -0.07, NS | 0.06, NS | 0.01, NS |

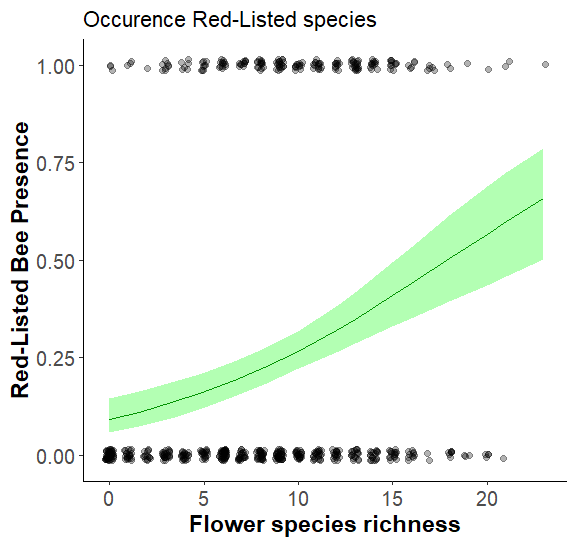

Figure D.14 The presence of at least one red listed species in a survey along a flower species richness gradient. Dark green line gives model predictions, the green area the 95% confidence interval. As red-listed species occurred in low abundance and species richness in each survey, a non-binary approach was not informative.

Appendix D.15: AIC values from models on Red-Listed bee presence with different polynomials (in rows) incorporating different landscape level factors or no landscape factor (columns). NNN stands for ‘NatuurNetwerk Nederland’, a network that includes green infrastructure and natural areas in the Netherlands. In grey the most parsimonious model (lowest AIC values).

|  | **Poly-nomial** | **NNN in 1km** | **NNN in 250m** | **Shortest distance to NNN** | **Agricultural area in 250m** | **Agricultural area in 1000m** | **Grassland area in 250m** | **Grassland area in 1000m** | **No landscape factor** |
| --- | --- | --- | --- | --- | --- | --- | --- | --- | --- |
| **Red-Listed Species Presence** | 1 | 577.46 | 585.49 | 590.44 | 588.51 | 588.77 | 587.28 | 585.49 | 587.28 |
|  | 2 | 578.54 | 586.27 | 590.98 | 588.77 | 589.18 | 587.77 | 585.91 | 587.78 |
|  | 3 | 580.41 | 588.23 | 592.98 | 590.74 | 591.14 | 589.73 | 587.82 | 589.74 |
|  | 4 | 582.41 | 590.23 | 594.92 | 592.74 | 593.14 | 591.73 | 589.81 | 591.74 |

Appendix D.16: Estimates and P-values and for different factors (in columns) from models on Red-Listed bee incorporating different landscape level factors or no landscape factor (rows). The final column gives the model marginal R^2^ value. NNN stands for ‘NatuurNetwerk Nederland’, a network that includes green infrastructure and natural areas in the Netherlands. In grey the most parsimonious model (lowest AIC value).

|  | **Flower Species Richness** | **Julian Date** | **Landscape** | **Marginal R^2^** |
| --- | --- | --- | --- | --- |
| **NNN in 1km** | 0.58, *** | -1.24, *** | 0.41, *** | 0.68 |
| **NNN in 250m** | 0.61, *** | -1.27, *** | 0.27, * | 0.65 |
| **Shortest distance to NNN** | 0.66, *** | -1.29, *** | -0.09, NS | 0.63 |
| **Agricultural area 250m** | 0.65, *** | -1.32, *** | -0.18, NS | 0.64 |
| **Agricultural area 1km** | 0.64, *** | -1.33, *** | -0.17, NS | 0.64 |
| **Grassland area 250m** | 0.65, *** | -1.31, *** | 0.22, NS | 0.65 |
| **Grassland area 1km** | 0.64, *** | -1.32, *** | 0.27, * | 0.65 |
