## Appendix E for "Phenology and flower species availability define wild bee communities on river embankments"

**Appendix E – Phenology**

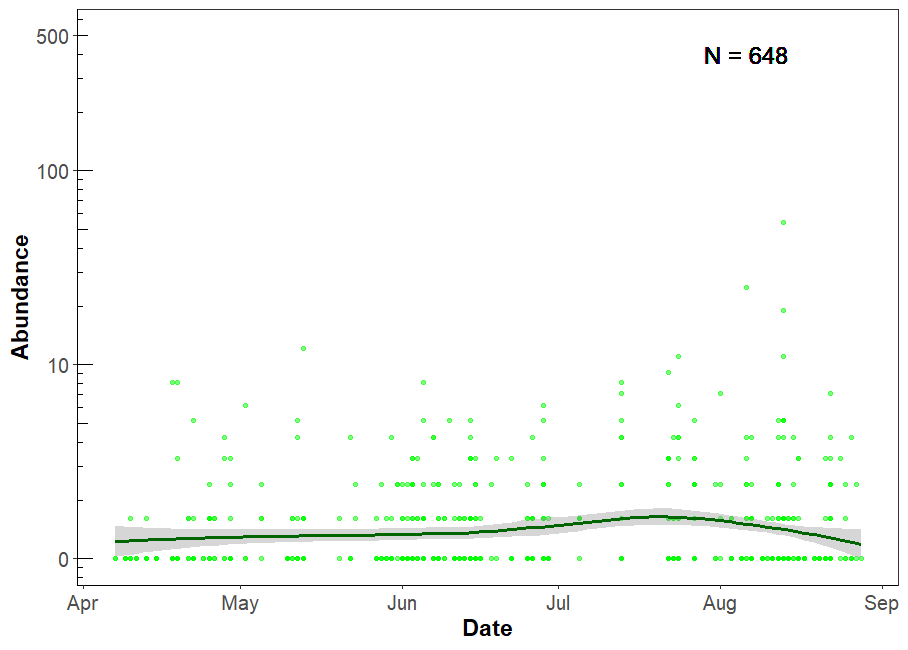

Appendix E.1: Encountered abundance of other genera per transect through the year on a log scale, along the Julian data. N gives the total number of individuals observed within each group, with each dot representing the within group number of observed individuals during a single transect count. We fitted a loess curve (dark green) with 95% confidence interval to aid interpretation.

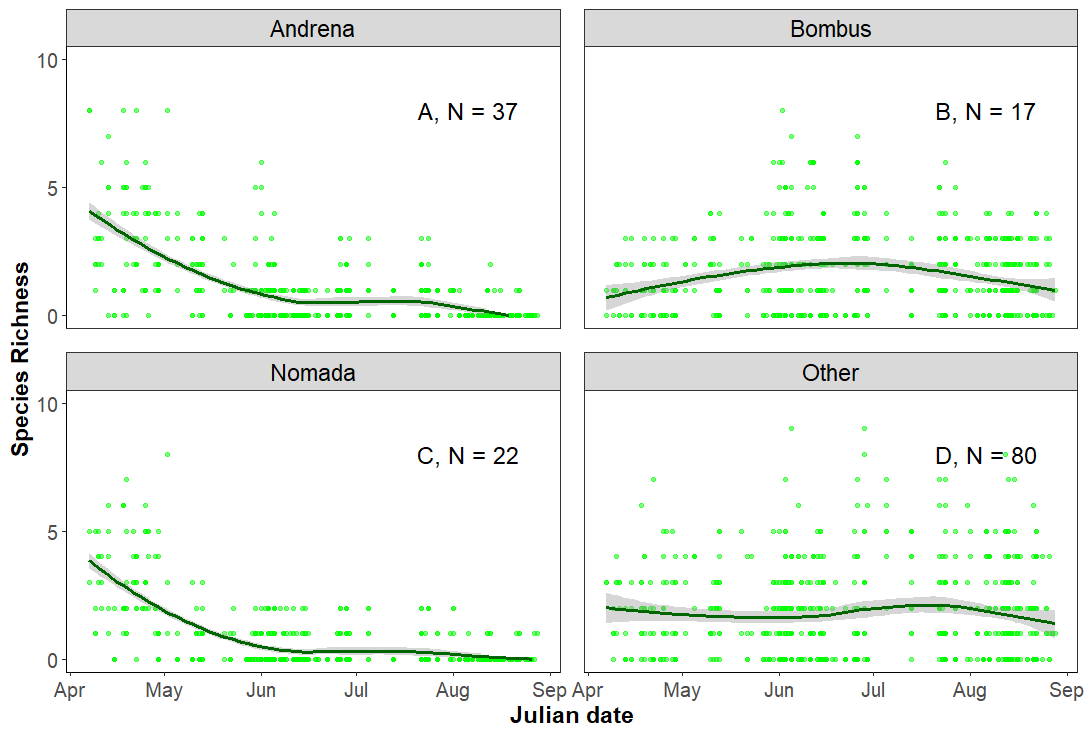

Appendix E.2: Species richness per transect within the three largest genera (Andrena (A), Bombus (B), Nomada (C)) and other genera (D) over the year along the Julian data. N gives the total number of species observed within each group, with each dot representing the within group species richness of observed individuals during a single transect count. We fitted a loess curve (dark green) with 95% confidence interval to aid interpretation.

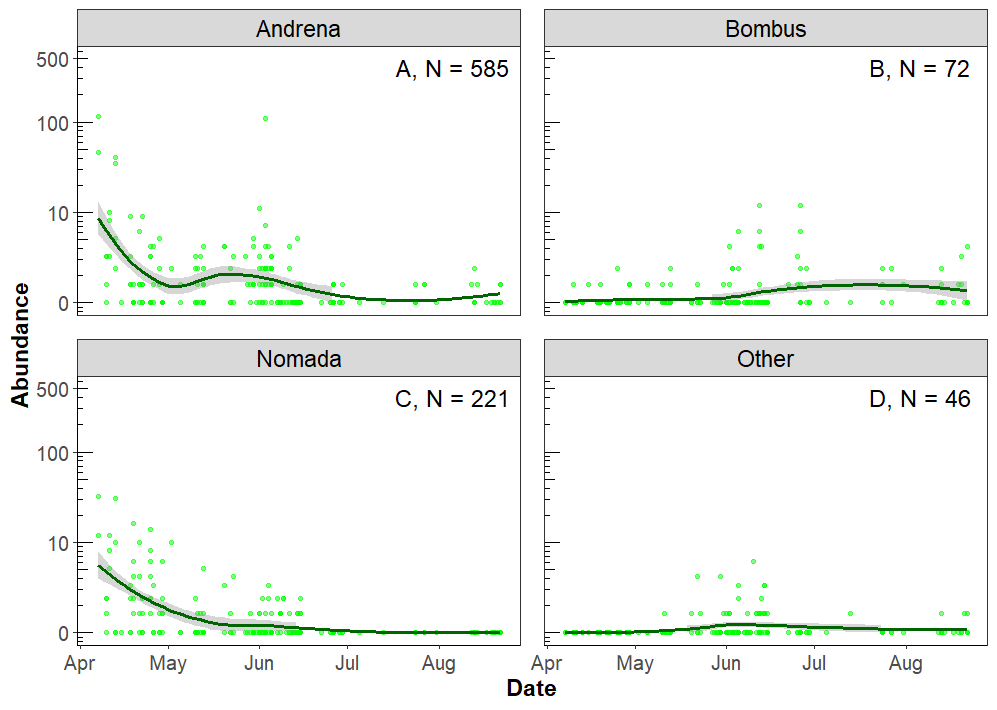

Appendix E.3: Encountered abundance of individuals of red-listed species within the three largest genera (Andrena (A), Bombus (B), Nomada (C)) and other genera (D) per transect over the year on a log scale, along the Julian data. N gives the total number of individuals observed within each group, with each dot representing the within group number of observed individuals during a single transect count. We fitted a loess curve (dark green) with 95% confidence interval to aid interpretation
