## Appendix F for "Phenology and flower species availability define wild bee communities on river embankments"

**Appendix F – Species specific relationships**

Appendix F.1: results of the presence-absence model per species from the core community. First column lists the species presence/absence was derived for, the second column its status in the Dutch Red-List for bees (Reemer, 2018) (VU = vulnerable, LC = least concern). The last three columns show the significance level of the flower species richness independent variable and model conditional and marginal Pseudo-R^2^.

| **Species** | **Red List Status** | **P-value** | **Pseudo R-squared – conditional** | **Pseudo R-squared - marginal** |
| --- | --- | --- | --- | --- |
| *Andrena dorsata* | LC | ** | 0.81 | 0.19 |
| *Andrena flavipes* | LC | *** | 0.66 | 0.56 |
| *Andrena gravida* | VU | NS | 0.71 | 0.24 |
| *Andrena haemorrhoa* | LC | ** | 0.73 | 0.55 |
| *Andrena labialis* | VU | * | 1.00 | 0.02 |
| *Andrena nitida* | LC | NS | 0.56 | 0.21 |
| *Andrena vaga* | LC | - | - | - |
| *Andrena wilkella* | VU | * | 0.99 | 0.08 |
| *Bombus hortorum* | LC | ** | 0.71 | 0.55 |
| *Bombus lapidarius* | LC | *** | 0.42 | 0.42 |
| *Bombus pascuorum* | LC | *** | 0.63 | 0.58 |
| *Bombus pratorum* | LC | ** | 0.80 | 0.18 |
| *Bombus terrestris/lucorum* | LC | *** | 0.48 | 0.42 |
| *Dasypoda hirtipes* | LC | *** | 0.69 | 0.28 |
| *Halictus scabiosae* | LC | ** | 0.78 | 0.45 |
| *Halictus tumulorum* | LC | *** | 0.57 | 0.20 |
| *Heriades truncorum* | LC | *** | 0.80 | 0.15 |
| *Hylaeus dilatatus* | LC | * | 0.58 | 0.58 |
| *Lasioglossum calceatum* | LC | *** | 0.47 | 0.33 |
| *Lasioglossum leucozonium* | LC | *** | 0.49 | 0.42 |
| *Lasioglossum malachurum* | LC | NS | 0.96 | 0.03 |
| *Lasioglossum morio* | LC | ** | 0.61 | 0.16 |
| *Lasioglossum pauxillum* | LC | *** | 0.66 | 0.39 |
| *Lasioglossum villosulum* | LC | ** | 0.79 | 0.21 |
| *Lasioglossum xanthopus* | LC | NS | 0.97 | 0.11 |
