## Appendix A for "Phenology and flower species availability define wild bee communities on river embankments"

**Appendix A – Breakpoint analysis**

Appendix A.1: description of flower abundance analysis
As flower abundance was measured in different, and not unifiable ways by the different observers, we analysed the relationship between bee abundance and diversity and flower abundance separately for each dataset. LS estimated flower coverage using the Tansley scale, CS counted all flowering plants within a 5x5m square and IR Counted all individual flowering units along their transect. Models were developed in similar ways as described in 2.3.3.

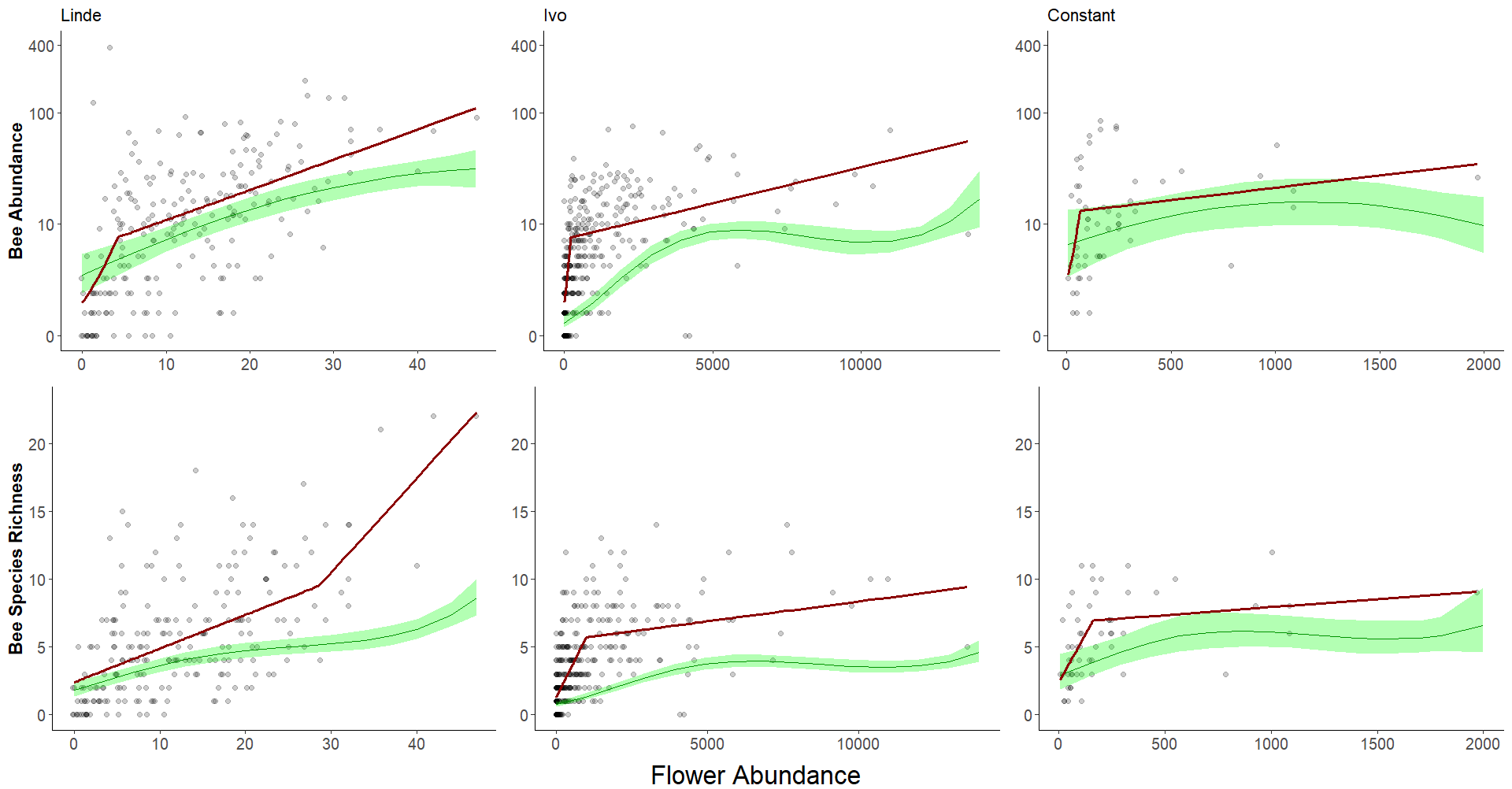

Appendix A.2: Correlations between all wild bee abundance (upper row) and diversity (lower row) and local flower abundance from the three different observer datasets. Red lines represent the outcomes of the breakpoint analysis on the linear relationships between the response variable (Y-axis) and Flower species richness.

Appendix A.3: Table showing the outcomes of the breakpoint analysis. The first column indicates the response variable, the second the estimated breakpoint, the third the slope before the breakpoint and the fourth the slope after the breakpoints. Outcomes are shown in Appendix 12.

| **Response variable** | **Breakpoint** | **Slope before** | **Slope after** |
| --- | --- | --- | --- |
| Abundance Linde | 4.25 | 0.38 | 0.06 |
| Abundance Ivo | 212 | 0.008 | 0.0002 |
| Abundance Constant | 66 | 0.024 | 0.005 |
| Diversity Linde | 28.67 | 0.25 | 0.45 |
| Diversity Ivo | 996.00 | 0.0045 | 0.0003 |
| Diversity Constant | 163.89 | 0.028 | 0.0011 |

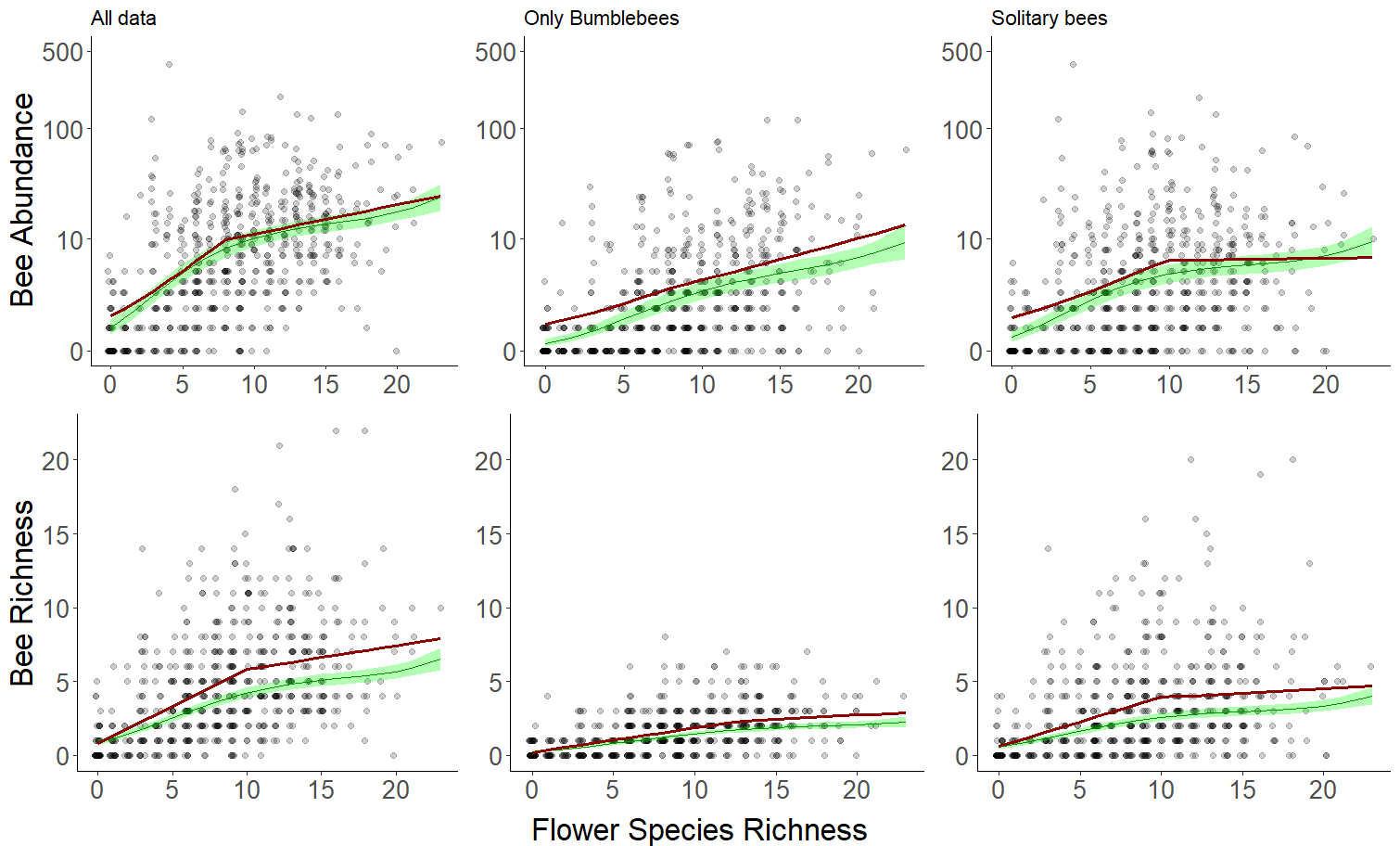

Appendix A.4: Correlations between abundance (upper row) and diversity (lower row) and local flower species richness from different subsets of the bee community; left including all available bee observation, middle for only bumblebees and right for non-Bombus bees. Red lines represent the outcomes of the breakpoint analysis on the linear relationships between the response variable (Y-axis) and Flower species richness.

Appendix A.5: Table showing the outcomes of the breakpoint analysis. The first column indicates the response variable, the second the estimated breakpoint, the third the slope before the breakpoint and the fourth the slope after the breakpoints. Outcomes are shown in Appendix 10.

| **Response variable** | **Breakpoint** | **Slope before** | **Slope after** |
| --- | --- | --- | --- |
| Abundance All Data | 7.88 | 0.23 | 0.06 |
| Abundance Only Bumblebees | 8.00 | 0.14 | 0.09 |
| Abundance Solitary Bees | 9.83 | 0.15 | 0.004 |
| Diversity All Bees | 10 | 0.50 | 0.16 |
| Diversity Bumblebees | 13 | 0.16 | 0.05 |
| Diversity Solitary Bees | 9.93 | 0.33 | 0.06 |
