## Appendix B for "Phenology and flower species availability define wild bee communities on river embankments"

**Appendix B – Data handling and data description**

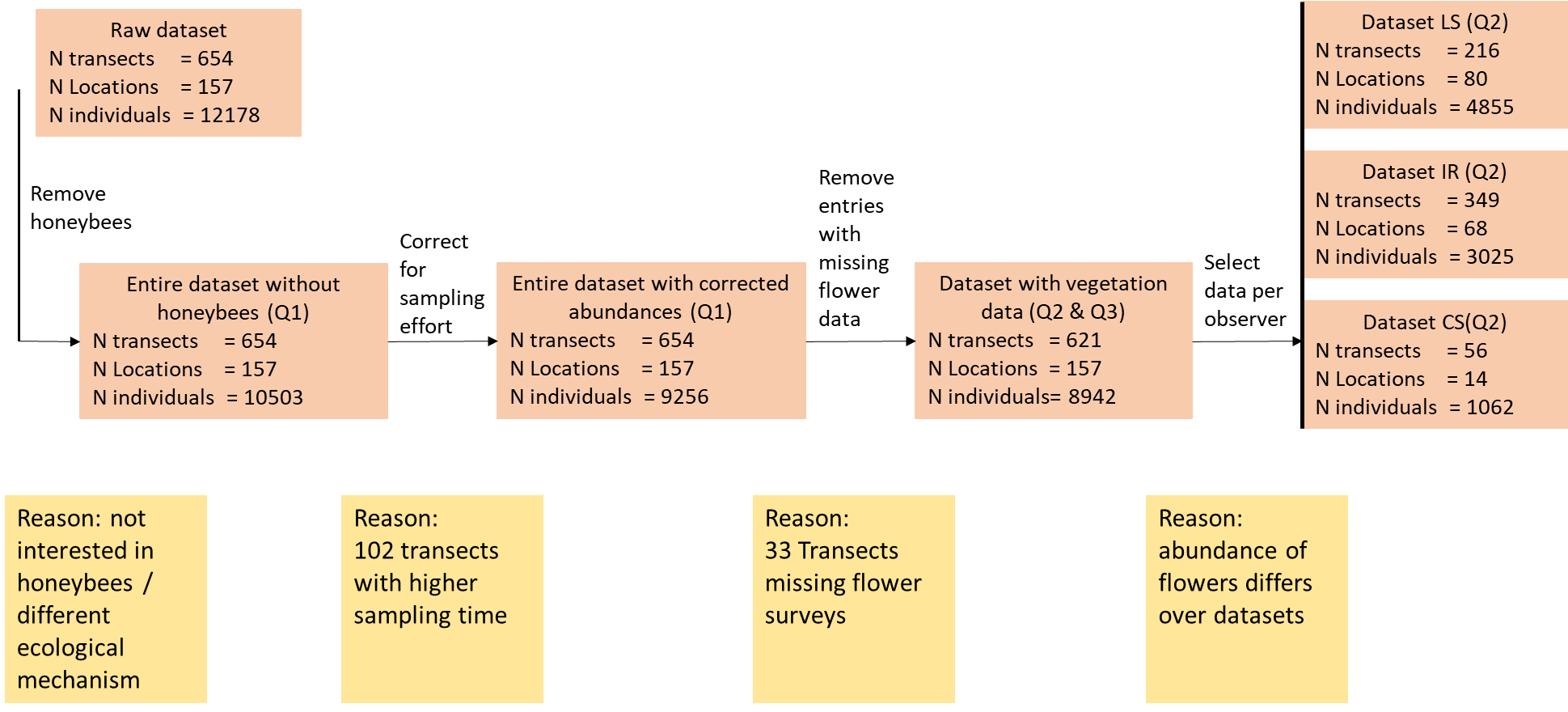
Appendix B.1: A flowchart showing the data selection process for answering our different research questions, indicated between brackets. A short reasoning is given as to why we made these selections.

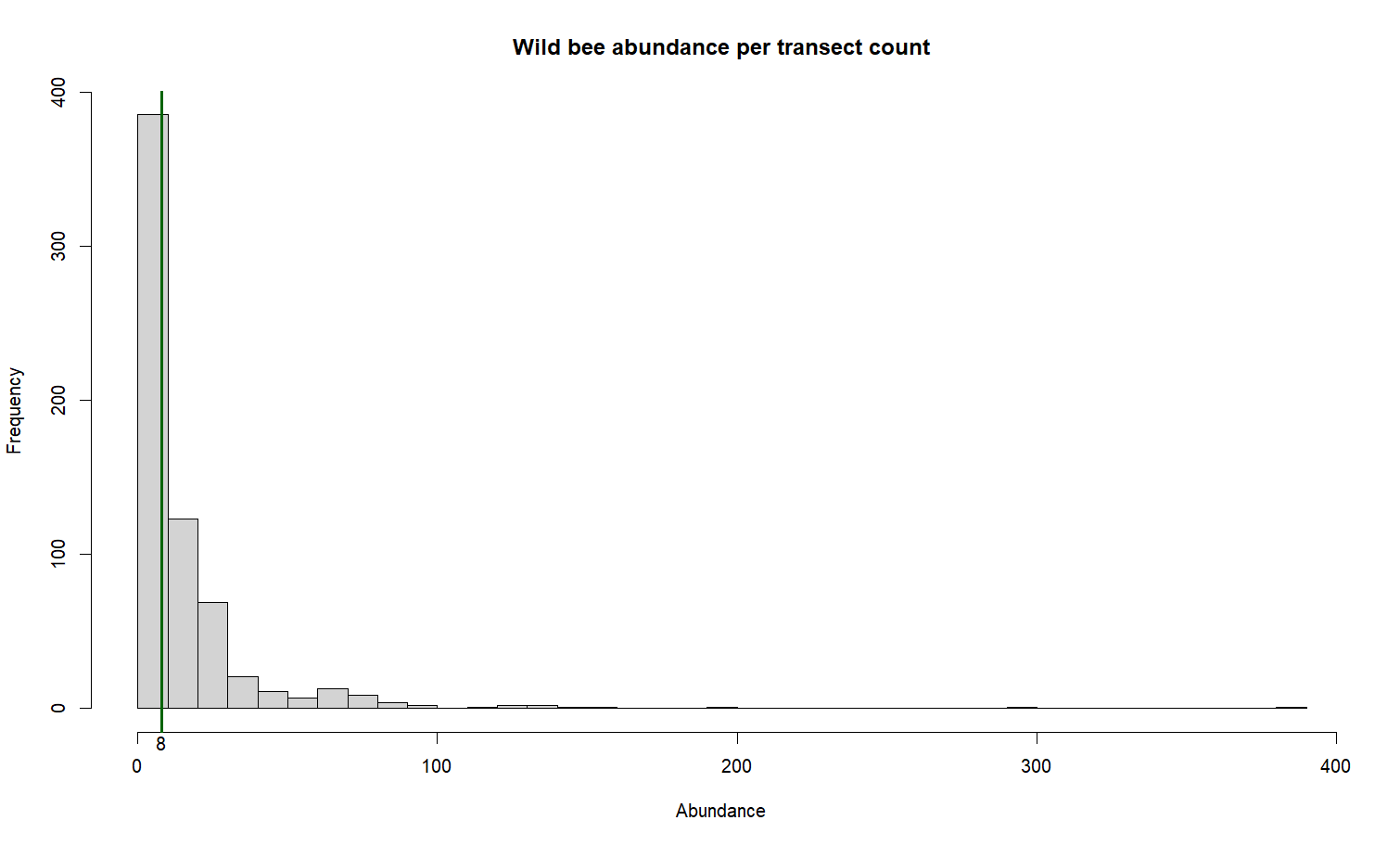

Appendix B.2: histogram showing the number of observed wild bee individuals per transect count. Green line indicates the median value.
